# Estimating rates and patterns of diversification with incomplete sampling: A case study in the rosids

**DOI:** 10.1101/749325

**Authors:** Miao Sun, Ryan A. Folk, Matthew A. Gitzendanner, Robert P. Guralnick, Pamela S. Soltis, Zhiduan Chen, Douglas E. Soltis

## Abstract

**Premise of the Study:** Recent advances in generating large-scale phylogenies enable broad-scale estimation of species diversification rates. These now-common approaches typically (1) are characterized by incomplete coverage without explicit sampling methodologies, and/or (2) sparse backbone representation, and usually rely on presumed phylogenetic placements to account for species without molecular data. Here we use an empirical example to examine effects of incomplete sampling on diversification estimation and provide constructive suggestions to ecologists and evolutionists based on those results.

**Methods:** We used a supermatrix for rosids, a large clade of angiosperms, and its well-sampled subclade Cucurbitaceae, as empirical case studies. We compared results using this large phylogeny with those based on a previously inferred, smaller supermatrix and on a synthetic tree resource with complete taxonomic coverage. Finally, we simulated random and representative taxon sampling and explored the impact of sampling on three commonly used methods, both parametric (RPANDA, BAMM) and semiparametric (DR).

**Key Results:** We find the impact of sampling on diversification estimates is idiosyncratic and often strong. As compared to full empirical sampling, representative and random sampling schemes either depress or exaggerate speciation rates depending on methods and sampling schemes. No method was entirely robust to poor sampling, but BAMM was least sensitive to moderate levels of missing taxa.

**Conclusions:** We (1) urge caution in use of summary backbone trees containing only higher-level taxa, (2) caution against uncritical modeling of missing taxa using taxonomic data for poorly sampled trees, and (3) stress the importance of explicit sampling methodologies in macroevolutionary studies.

## I. Introduction

With recent advances in generating very large phylogenetic trees (e.g., Smith and O’Meara, 2012; Stamatakis, 2014; Hinchliff et al., 2015; Nguyen et al., 2015; Smith and Brown, 2018; Eiserhardt et al., 2018), and in analytical methods (Nee et al., 1994a,b; Pybus and Harvey, 200; Paradis et al., 2004; Alfaro et al., 2009; Stadler, 2011; Pennell et al., 2014; Rabosky, 2014; Morlon et al., 2016; Höhna et al., 2016), assessing macroevolutionary patterns for globally distributed clades with available biodiversity information has become common (e.g., Jetz et al., 2012; Morlon, 2014; Scholl and Wiens, 2016; Magallón et al., 2018; Rabosky et al., 2018; Upham et al., 2019). Analyses of diversification rates have shed light on potential drivers of diversity gradients across wide phylogenetic and geographic scales (Jetz et al., 2012; Rabosky et al., 2018; Landis et al., 2018). However, inferring diversification processes based solely on extant species phylogenies is very challenging (Etienne et al., 2011; Didier et al., 2017; Sauquet and Magallón, 2018; Mitchell et al., 2019), and the accuracy of these methods is an area of intensive research and heated controversy (O’Meara and Beaulieu, 2016; Moore et al., 2016; Rabosky et al., 2017; Meyer et al., 2018; Rabosky, 2018). Many contemporary analytical workflows for studying diversification have seen little vetting to date with empirical datasets (but see Title and Rabosky, 2017), and much remains to be explored about the response of diversification methods to missing and biased species sampling (Sauquet and Magallón, 2018).

On the empirical side, incomplete sampling of molecular phylogenetic data for many clades represents a long-standing constraint on assembling datasets to adequately explore large-scale macroevolutionary questions (e.g., Linder et al., 2005; Cusimano et al., 2012; Thomas et al., 2013). Diversification models generally have no information from which to draw inferences other than branching order and branch length among extant species, both of which can be dramatically affected by (1) absolute taxon coverage (FitzJohn et al., 2009; Dvies et al., 2013; Title and Rabosky, 2017; Revell, 2018; Burin et al., 2018; Rabosky, 2018) and (2) sampling method at a given level of taxon coverage (Höhna et al., 2011; Höhna, 2014; Cusimano et al., 2012). Hence, not only absolute taxon coverage, but also potential bias in this coverage, is important in interpreting diversification results, yet the identification and use of explicit sampling strategies remains uncommon in the field (O’Meara et al., 2016). Inclusion of data representing all extant lineages with molecular data from resources like GenBank, without an explicit sampling methodology, is perhaps the most common analytical strategy (e.g., Jetz et al., 2012; Zanne et al., 2014; Upham et al., 2019; but see, e.g., Magallón et al., 2018; O’Meara et al., 2016). A second commonly used approach is taxonomically representative sampling, including family-level or genus-level backbone trees (e.g., Magallón et al., 2018), which preferentially samples species to represent deep phylogenetic divergences to the exclusion of recent divergences. Representative sampling is the community standard for molecular phylogenetic studies, meaning that databases such as GenBank implicitly contain representative bias (reviewed in Cusimano et al., 2012; Höhna, 2014; O’Meara et al., 2016; Sauquet and Magallón, 2018). Finally, random sampling procedures that sample extant species with equal probability are perhaps the least frequently used (although this approach corresponds best to common model assumptions; see O’Meara et al., 2016).

Most current diversification approaches are able to model incomplete sampling; a variety of such methods is widely used in recent diversification studies (as a small sample across taxa: Jetz et al., 2012; Rabosky et al., 2018; Magallón et al., 2018). Methods for accounting for missing taxa make strong assumptions about the structure of missing species, typically assuming they are randomly missing, an assumption not matched in many empirical datasets (Höhna et al., 2011; Cusimano et al., 2012; Thomas et al., 2013; Revell, 2018), and the impact of alternative sampling approaches is not clear. An additional poorly understood area is the impact of methods for incorporating described taxonomic diversity for which molecular phylogenetic data are unavailable. The increased availability of very large synthetic phylogenetic products with backbone taxonomy such as the Open Tree of Life (Hinchliff et al., 2015), as well as probabilistic methods for inserting backbone taxonomic information (e.g., polytomy resolver, Kuhn et al., 2011; PASTIS, Thomas et al., 2013; TACT, Rabosky et al., 2018), creates opportunities for very large analyses with complete sampling of known diversity. However, while these methods are often used (e.g., Jetz et al., 2012; Rabosky et al., 2018; see review by Rabosky, 2015), the properties of diversification inference with contemporary methods using such backbone taxonomies remain poorly understood.

Here we use the rosid clade in the flowering plants as a test case to explore how different sampling schemes influence the estimation of diversification with empirical data. Rosids (*Rosidae*; Cantino et al., 2007; Wang et al., 2009; APG IV, 2016) have great potential for understanding the evolution and diversification of angiosperms, considering their enormous species richness (90,000–120,000 species, representing around 25% of all angiosperms; Govaerts, 2001; Hinchliff et al., 2015; Folk et al., 2018). The clade, containing such globally important families as grapes, legumes, oaks and beeches, squash and melons, and mustards (respectively, Vitaceae, Fabaceae, Fagaceae, Cucurbitaceae, and Brassicaceae), originated in the early to late Cretaceous (115 to 93 million years ago, hereafter Myr), followed by rapid diversification in perhaps as little as 4 to 5 million years to yield the crown groups of fabids (112 to 91 Myr) and malvids (109 to 83 Myr; Wang et al., 2009; Bell et al., 2010; Magallón et al., 2015). The rise of the rosids yielded today’s forests, which largely remain dominated by rosid species. The advent of these forests spurred diversification in many other lineages of life (e.g., ants: Moreau et al., 2006; Moreau and Bell, 2013; amphibians: Roelants et al., 2007; mammals: Bininda-Emonds et al., 2007; fungi: Hibbett and Matheny, 2009; liverworts: Feldberg et al., 2014; ferns: Watkins and Cardelús, 2012; Testo and Sundue, 2016). However, biodiversity knowledge in the rosids remains limited, with perhaps only 23% of species having usable molecular data for phylogenetics (i.e., not repetitive DNA and other non-conserved markers; Folk et al., 2018). Sampling is likewise biased; species coverage is highly uneven, with economically important groups like the legume and beech orders (Fabales, Fagales) overrepresented compared to important but less familiar tropical groups like Malpighiales (Folk et al., 2018).

Despite previous efforts assessing the impact of incomplete sampling (e.g., Cusimano et al., 2012; Höhna, 2014; Title and Rabosky, 2017), much remains unknown about how incomplete and biased taxon sampling approaches impact diversification estimates, particularly with empirical supermatrix data. Additionally, much of the methodological literature cited above does not include use of the most recent methods now widely used in the community. While offering limited power to generate biological insight about the diversification process, incomplete taxon coverage in the rosids is an opportunity to characterize the robustness of contemporary methods with an empirical dataset. We used a recently constructed, 5-locus, 19,700-taxon matrix for rosids (molecular data only; hereafter, *20k-tip tree*; Sun et al., 2019) to compare with a previously published 4-locus, 8,855-taxon rosid phylogeny (molecular data only; hereafter, *9k-tip tree*; Sun et al., 2016) as well as the rosid clade extracted from Open Tree of Life (hereafter OpenTree) with complete species sampling (molecular data and backbone taxonomic data; hereafter, *100k-tip tree*; Hinchliff et al., 2015; Smith and Brown, 2018). We explored results generated using these phylogenies from a suite of commonly used diversification approaches, comprising two parametric methods (RPANDA, Morlon et al., 2016; BAMM, Rabosky, 2014) and one semi-parametric method (the DR statistic, Jetz et al., 2012). We examined both variation in empirical sampling patterns in major rosid clades and a series of sampling perturbations to simulate random and representative sampling methods. Using the workflow summarized in Fig. 1, we document a remarkably complex impact of taxon sampling on inference of macroevolutionary patterns. We focused on the following questions: (1) Do commonly used contemporary methods differ in their robustness to poor overall sampling? (2) Do datasets generated by random and representative sampling strategies result in different diversification inferences? (3) Does adding backbone taxonomic information actually improve diversification inference?

**Fig. 1.**
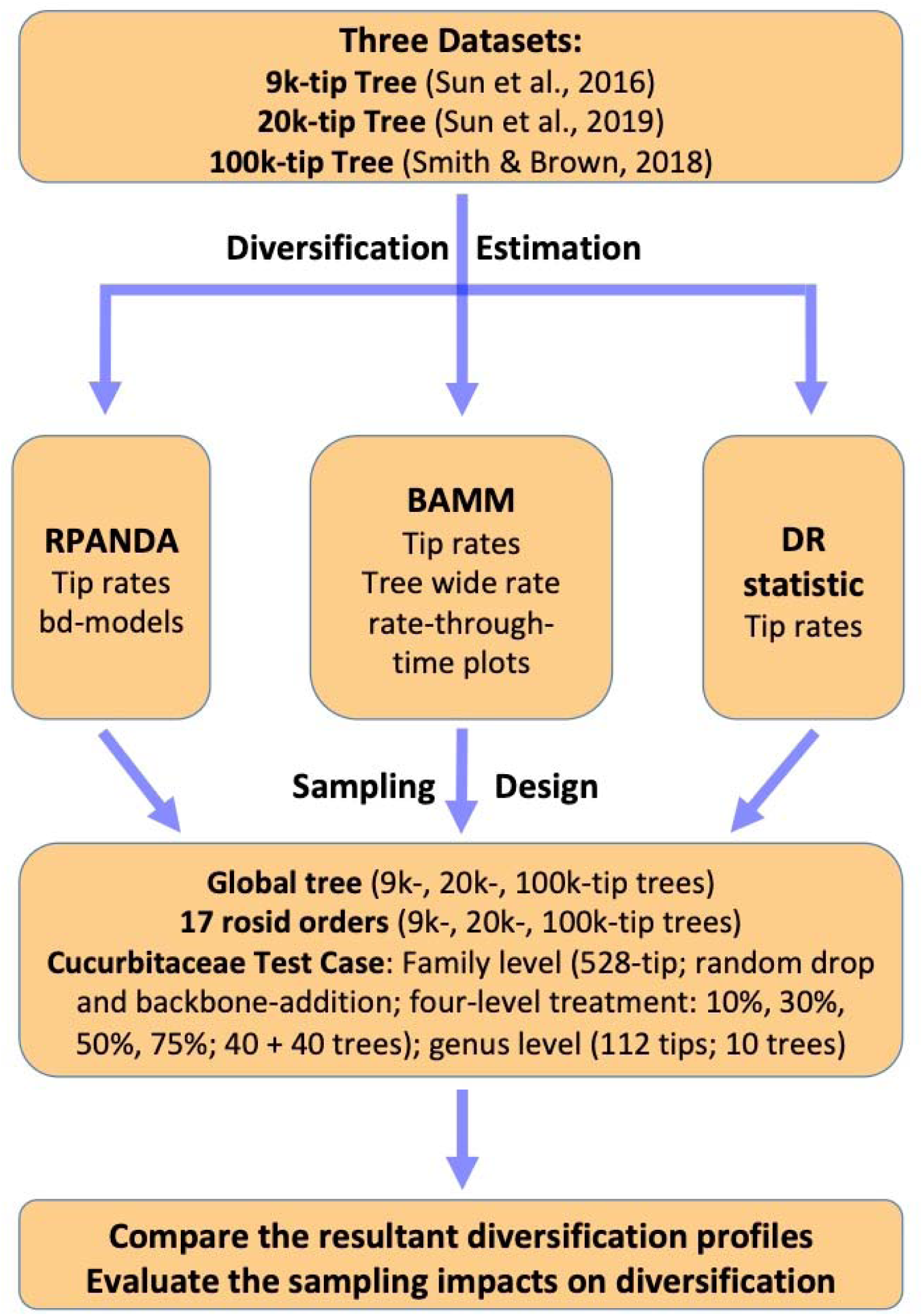
Workflow employed for empirical data and simulations in this study. Abbreviation notes: 9k-tip tree = 4-gene, 8,855-species rosid tree from Sun et al. (2016); 20k-tip tree = 5-locus, 19,740-species rosid tree from Sun et al. (2019); 100k-tip tree = 106,910-species tree extracted from OpenTree (Smith and Brown, 2018); bd-models = nine birth-death models from RPANDA (see Appendix S1.1). Tree-wide rate means speciation rate averaged throughout the tree.

## II. Materials and Methods

### The 9k-tip tree

This tree is the 4-gene tree of Sun et al. (2016) based on three chloroplast loci (*atpB*, *rbcL*, and *matK*) and one mitochondrial locus (*matR*). Details of its construction can be found in Sun et al. (2016). The data set consists of 8,855 ingroup species with 59.26% missing data and is largely congruent with other phylogenetic results for rosids (e.g., Wang et al., 2009; Soltis et al., 2011; Ruhfel et al., 2014; Gitzendanner et al., 2018).

### The 20k-tip tree

The 20k-tip tree was built by adding the nuclear ITS locus to the four genes in the 4-gene matrix of Sun et al. (2016), resulting in a 5-locus matrix with 19,740 ingroup species (135 families and 17 orders) and 70.55% missing data (See Sun et al., 2019). All families are monophyletic, and this phylogeny is also largely congruent with other inferences of rosid phylogeny (e.g., Wang et al., 2009; Soltis et al., 2011; Sun et al., 2016; Gitzendanner et al., 2018).

### The 100k-tip tree

We also assembled a complete species-level tree for all named rosid species using OpenTree. We pruned the rosid clade from a recent phylogeny dating all seed plants in OpenTree (see details in Smith and Brown, 2018; https://github.com/FePhyFoFum/big_seed_plant_trees/releases; file ALLOTB.tre), removed non-species designations as above, and smoothed the branch lengths after pruning. These steps were completed via functions from Phyx (Brown et al., 2018) and scripts from OpenTree PY Toys (https://github.com/blackrim/opentree_pytoys). The final cleaned tree contained 106,910 tips.

Divergence time analyses for these three trees (9k-, 20k-, and 100k-tip trees) have already been conducted previously (see details from Sun et al., 2019, and Smith and Brown, 2018, respectively; Fig. 2). Briefly, Sun et al. (2019) used treePL with 59 fossil constraints for the 9k-tip (Sun et al., 2016) and the 20k-tip phylogenies; likewise, Smith and Brown (2018) also used treePL with 590 constraints extracted from Magallón et al. (2015).

**Fig. 2.**
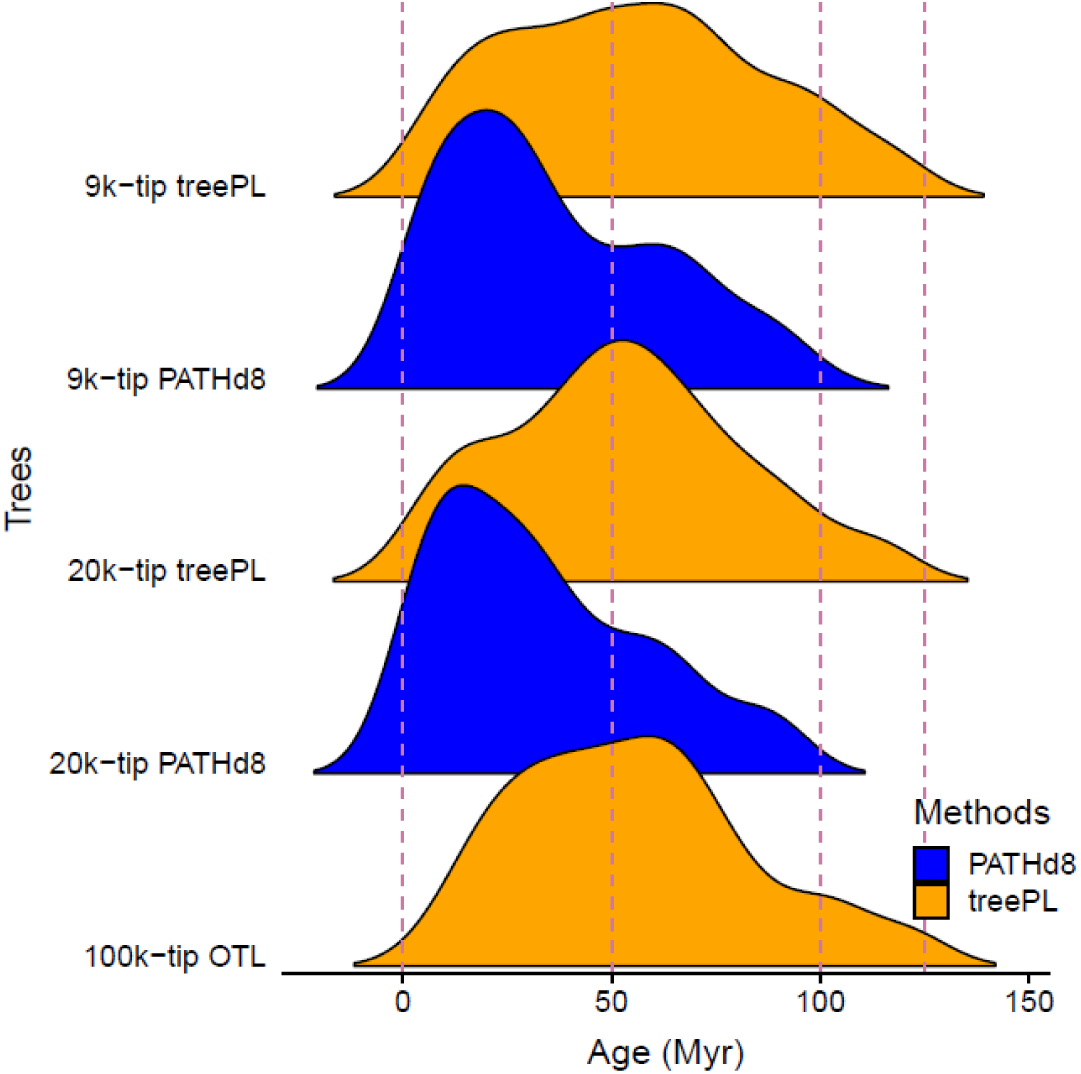
Age distribution of crown ages for clades extracted from the 9k-, 20k-, and 100k-tip trees. The two dating methods used, treePL and PATHd8, are shown in orange and blue, respectively. The two methods resulted in substantially different date scaling; for the treePL trees used in this study, the probability density distributions of clade dates were very similar across very different sampling levels.

### Diversification Analyses and Comparisons

To understand the impact of sampling strategies, we first used trends in empirical sampling across the three trees to investigate the correlation between sampling and inferred diversification. We compared patterns for both the overall trees and for the 17 orders (each monophyletic) of the rosid clade (APG IV, 2016), the species-level sampling of which differs by up to 8-fold among the trees. We applied three widely used contemporary methods: RPANDA (Morlon et al., 2016), BAMM (Rabosky, 2014), and the DR statistic (Jetz et al., 2012; for implementation details, see below). To generate comparable metrics across methods, we focused on the diversification rate of present-day lineages (that is, speciation rate at time zero or “tip rate”), a metric that is commonly used and is comparable across all of the methods employed (see Title and Rabosky, 2019). We used both global tip speciation rates (that is, speciation rates estimated at present, averaging across species; RPANDA, BAMM) and distributions of rates for individual contemporary species (=“tip rates”; BAMM, DR). For BAMM, we additionally examined speciation rates throughout the timeline of the phylogeny, using both averages across the entire tree (hereafter, tree-wide speciation rates) and rate-through-time plots.

### Sampling Treatments: Cucurbitaceae Test Case

To examine diversification patterns further by generating known sampling patterns, we used the best-sampled rosid family (Cucurbitaceae; approximately 64% sampling following *Flora of North America* [Nesom, 2015] and *Flora of China* [Lu et al., 2011]).

*Sampling treatments—*We extracted the Cucurbitaceae clade (a subset of 528 tips) from the 20k-tip tree to maximize species representation with molecular data alone. We simulated both random and representative sampling schemes, the former with and without backbone taxonomies. We (1) simulated randomly missing species by generating trees, randomly dropping extant species at four sampling levels (10%, 30%, 50%, and 75% of sampled species), with ten replicates for each sampling treatment. We then (2) simulated randomly missing species that are added in via backbone taxonomies (hereafter, “backbone-addition”) via randomly dropping extant species at four sampling levels (10%, 30%, 50%, and 75% of sampled species) and then adding them back to the phylogeny by attaching them to the most recent common ancestor (MRCA) of the genus, with the tip branch length extended to the present, similar to the method of OpenTree. If there were not at least two species of a genus sampled to generate a genus node, the missing taxon was attached to the root of the tree (i.e., it was assignable to the family Cucurbitaceae but not to any sampled genus node). These steps were done in 10 replicates with OpenTree PY Toys (https://github.com/blackrim/opentree_pytoys). Finally, to simulate representative sampling, we (3) pruned this tree to a genus-level phylogeny by randomly selecting one species in each genus in ten replicates. Across these scenarios we repeated the diversification methods for empirical trees (above) on these replicate trees.

### Diversification methods

We used RPANDA v1.4 (Morlon et al., 2016), a likelihood method, to fit nine diversification models representing constant, linear, and exponential time-dependent pure-birth and birth-death models (Morlon et al., 2014; Appendix S1.1, see Supplemental Data with this article). The best model was chosen individually across all empirical datasets, and simulated replicates and parameters presented are always from the individual best model. We accounted for incomplete sampling in each analysis to test whether this is adequately modeled by RPANDA, basing the sampling ratio on the total species number in the Open Tree Taxonomy (“OTT”) database (Table 1). We extracted the speciation rate parameter at present for downstream analyses as a metric comparable to commonly used per-species “tip rates” derived below from BAMM and DR. This quantity represents global speciation rates estimated for extant taxa and hereafter will be denoted “global tip speciation rate”.

**Table 1.** Ordinal-level summary sampling table for the 9k- and 20k-tip rosid sampling compared to the rosid clade of the Open Tree Taxonomy (“OTT”) database v. 3.0 (https://devtree.opentreeoflife.org/about/taxonomy-version/ott3.0; Hinchliff et al., 2015) and matching taxon names between these data sets. Orders follow APG IV (2016). A summary table at the family level for the 20k-tip tree is available in Sun et al. (2019).

| **Order** | **9k-tip Tree** |  | **20k-tip Tree** |  |
| --- | --- | --- | --- | --- |
|  | **Matched**<br>**genus %** | **Matched**<br>**species %** | **Matched**<br>**genus %** | **Matched**<br>**species %** |
| Brassicales | 36.85% | 7.49% | 71.12% | 28.50% |
| Celastrales | 59.45% | 13.34% | 61.26% | 18.15% |
| Crossosomatales | 92.85% | 29.26% | 92.86% | 29.27% |
| Cucurbitales | 85.71% | 13.93% | 87.97% | 26.60% |
| Fabales | 66.66% | 8.25% | 76.04% | 21.95% |
| Fagales | 44.59% | 10.91% | 48.65% | 21.92% |
| Geraniales | 60.00% | 12.16% | 75.00% | 30.67% |
| Huerteales | 100.00% | 23.33% | 100.00% | 23.33% |
| Malpighiales | 64.98% | 8.33% | 65.77% | 17.37% |
| Malvales | 54.81% | 9.72% | 62.96% | 16.54% |
| Myrtales | 48.21% | 4.05% | 54.11% | 8.28% |
| Oxalidales | 59.42% | 4.21% | 62.32% | 8.25% |
| Picramniales | 66.66% | 8.77% | 66.67% | 8.77% |
| Rosales | 54.03% | 3.38% | 60.45% | 8.22% |
| Sapindales | 56.98% | 11.21% | 62.07% | 18.34% |
| Vitales | 60.00% | 3.63% | 60.00% | 9.52% |
| Zygophyllales | 62.96% | 10.58% | 66.67% | 17.65% |
| Total | 57.80% | 7.28% | 66.34% | 16.25% |

We used BAMM v2.5.0 (Rabosky, 2014), a Bayesian approach, to estimate tip speciation rates as with RPANDA (above). We also used BAMM to explore non-contemporary speciation rates, examining both tree-wide speciation rates (that is, speciation rates averaged across all tree timeframes including the present) and rate-through-time plots (that is, speciation rates averaged in temporal windows, Appendix S1.2). We also accounted for incomplete sampling in BAMM, parameterizing this identically to RPANDA (above).

As an additional examination of common practices, we used BAMM to explore the impact of a global sampling probability (one missing species proportion imposed as the parameter for the entire tree) and species-specific sampling probabilities (missing species parameters for arbitrarily defined clades, often named taxa) on diversification rates implemented in BAMM. We confirmed convergence of the MCMC chains and effective sample sizes >200 for the number of both shifts and log likelihoods (Appendix S2.1), after discarding 10% burn-in. The exception was in order-level BAMM analyses for the 100k-tip tree, for which 6 orders (Brassicales, Fabales, Malpighiales, Myrtales, Rosales, and Sapindales) could not reach suitable effective sample sizes despite runs in some cases exceeding 400 million generations; in these cases we imposed a 90% burn-in to ensure adequate convergence and reduce downstream computational time. We present results from these orders for comparison; results were qualitatively similar to other orders in the 100k-tip tree (see Results).

Lastly, we employed the DR statistic (Jetz et al., 2012), one of the most widely used semiparametric approaches to diversification estimation. The DR statistic quantifies the “splitting rate” from each extant species to the tree root as a model-free estimate of diversification rate. Methods followed those described in Jetz et al. (2012) and Harvey et al. (2016). There is no straightforward way to model incomplete sampling with the DR statistic (but see Rabosky et al., 2018); aside from calculating DR for our 100k-tip synthetic tree, we did not account for missing taxa in order to represent the most typical way in which this statistic has been used. For BAMM, it was impossible to achieve convergence in the global 20k-tip and 100k-tip trees, so we only ran this method on the 17 rosid orders (clades recognized in APG IV, 2016); global tree results were generated only for DR and RPANDA.

## III. Results

### Diversification Analyses

#### Empirical diversification patterns

*RPANDA—*Both the 9k-tip and 20k-tip trees favored a birth-death model with speciation and extinction rates varying exponentially with time; the optimal model for the 100k-tip tree was a pure birth model with linear speciation rate with respect to time (Appendix S1.1; Appendix S2.2). The tip speciation rate was highest for the 9k-tip tree (1.3905 Myr^-1^) with similarly high results from the 20k-tip tree (1.3058 Myr^-1^); estimated rates for the 100k-tip tree were much lower (0.0446 Myr^-1^; Fig 3a).

**Fig. 3.**
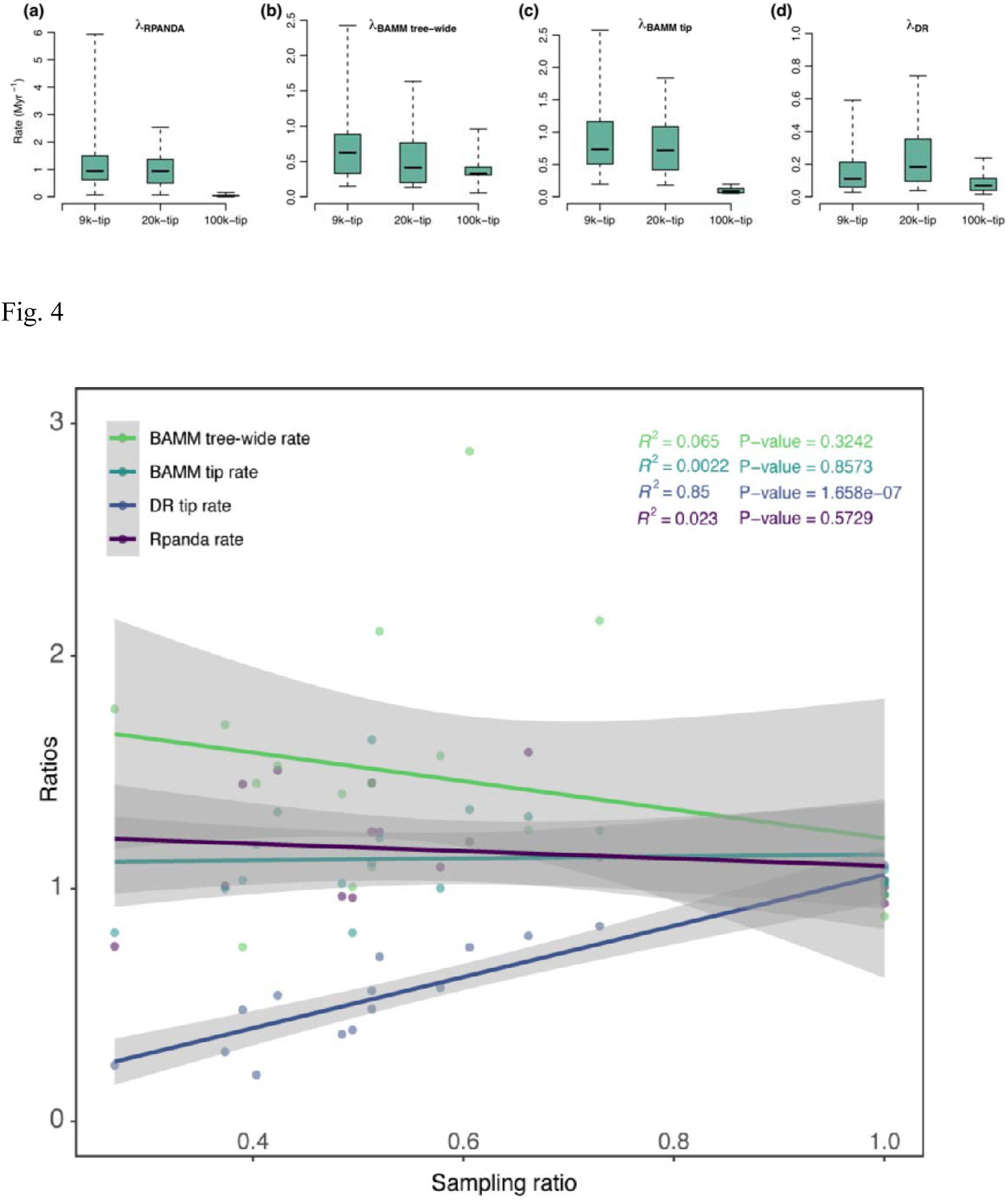
Tip speciation rate boxplots (here denoted λ_0_) for RPANDA, BAMM (including tip speciation rates and tree-wide speciation rates), and DR, across the three empirical datasets, 9k-tip tree, 20k-tip tree, and 100k-tip tree. The boxes and whiskers represent the 0.25–0.75 and the 0.05–0.95 quantile ranges, respectively.

*BAMM—*The values of both mean tip speciation rates and mean tree-wide speciation rates for the 9k-tip tree (1.1527 Myr^-1^ and 0.7829 Myr^-1^, respectively) are higher than those from both the 20k-tip tree (1.0731 Myr^-1^ and 0.5601 Myr^-1^; Appendix S2.1; Fig. 3) and 100k-tip tree (0.1136 Myr^-1^ and 0.3914 Myr^-1^; Appendix S2.1; Fig. 3b-c). Among the 17 orders, both the tip and tree-wide speciation rates from the 9k-tip tree are likewise generally slightly higher than the 20k-tip tree and much higher than the 100k-tip tree (Appendix S2.1; Fig. 3bc).

*DR—*On average, DR tip rates estimated from the 20k-tip tree yielded the highest value (0.4644 Myr^-1^), the 9k-tip tree was intermediate at 0.1889 Myr^-1^, while the 100k-tip tree yielded the lowest (0.0902 Myr^-1^; Appendix S2.3; Fig. 3d). As with the previous methods, this overall scaling was also generally true across the 17 orders (Appendix S2.3).

#### Sampling and diversification among rosid orders

RPANDA and BAMM showed a negative relationship between sampling ratio and estimated rates across the empirical data for the 17 rosid orders (that is, orders with less sampling effort had greater estimated speciation rates). However, this correlation was not significant (cf. Fig. 4). The DR method, however, showed a strong positive correlation (*p* = 1.658e-07) between sampling ratios and estimated rates, meaning that decreasing sampling effort predicts lower estimated speciation rates using this method (Fig. 4).

**Fig. 4.** Correlation between sampling effort and speciation rates among the 17 rosid orders from 9k- and 20k-tip trees. The X axis is the ratio of sampling percentages; the Y axis is the ratio of speciation rates (9k-tip/20k-tip in both cases; values closer to one indicate values closer to the more fully sampled 20k-tip tree); each dot represents a single rosid order. The *R^2^* and *p-*values are color-coded following the legend colors. Gray plot zones indicate curve 95% confidence intervals. Only the DR statistic showed a significant positive relationship between sampling percentage and diversification rate; for other methods, the rosid orders do not show a significant relationship between sampling effort and estimated speciation rate.

Rate-through-time curves across all orders showed strong differences among the three trees (Appendix S3). The 9k- and 20k-tip trees were most similar across analyses; however, the improved sampling of the 20k-tip tree allowed for the detection of recent bursts within the last 15 million years in several orders that were not inferred in the 9k-tip tree (e.g., Brassicales, Cucurbitales, Fabales, Malpighiales, Vitales; Appendix S3). The difference between the 100k-tip tree and the 9k- and 20k-tip trees was more substantial. In the 100k-tip tree, with the exception of Huerteales, all order analyses detected early bursts of speciation rate not found in other trees, with lower estimated tip rates (here, at time 0) than the 9k- and 20k-tip trees (also see Fig. 3c).

#### Cucurbitaceae test case—Random sampling simulation

*RPANDA—*With random sampling, the estimated global tip speciation rate increased with decreasing sampling effort, ranging about 1.5 fold from 0.4687 Myr^-1^ (10% random drop) to 0.7263 Myr^-1^ (75% random drop; Fig. 5a; Appendix S2.4). The 75% random-drop treatment was significantly higher in tip speciation rate than all other treatments; no other treatment comparisons were significantly different (Tukey HSD; see Appendix S2.5).

**Fig. 5.**
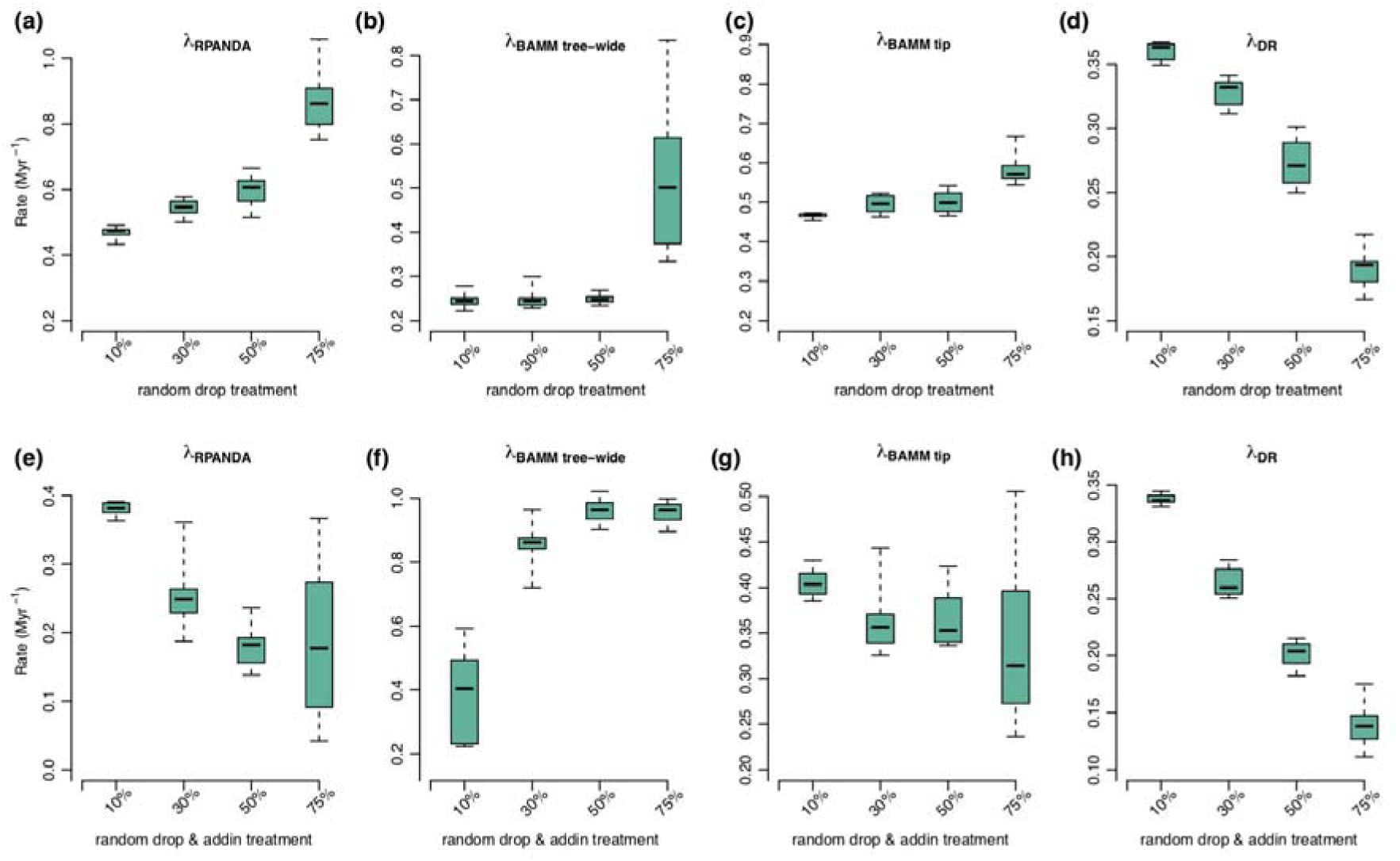
Sampling simulation boxplots with four treatments and three different rate metrics using the Cucurbitaceae tree. Contemporary speciation rates (λ) estimated by RPANDA (λ_RPANDA_), BAMM (speciation rate: λ_BAMM tree-wide_; and tip rate: λ_BAMM tip_), and DR (λ_DR_). The (a-d) panels correspond to the random sampling simulations and (e-f) panels to the random sampling simulations with backbone-addition.

*BAMM—*As with RPANDA, higher estimated mean tip speciation rates and tree-wide speciation rates were both associated with decreasing sampling effort under random sampling, ranging from 0.4658 Myr^-1^ to 0.6508 Myr^-1^ for mean tip speciation rates and from 0.2466 Myr^-1^ (10% randomly dropped) to 0.5261 Myr^-1^ (75% randomly dropped) for mean tree-wide speciation rates (Fig. 5b,c; see Appendix S2.4). These rates were statistically identical for all treatments except the 75% random-drop treatment (Tukey HSD; see Appendix S2.5).

Rate-through-time plots from the trees show a similar pattern (Fig. 6) to those observed for tip speciation rates. All of the sampling treatments tend to agree in rate magnitude and curve shape with the complete tree except for the 75% random drop treatment; in this treatment the overall speciation rates are higher at all timeframes, and the curves tend to be flattened and linearized, with few of the complex details apparent with greater sampling (Fig. 6).

**Fig. 6.**
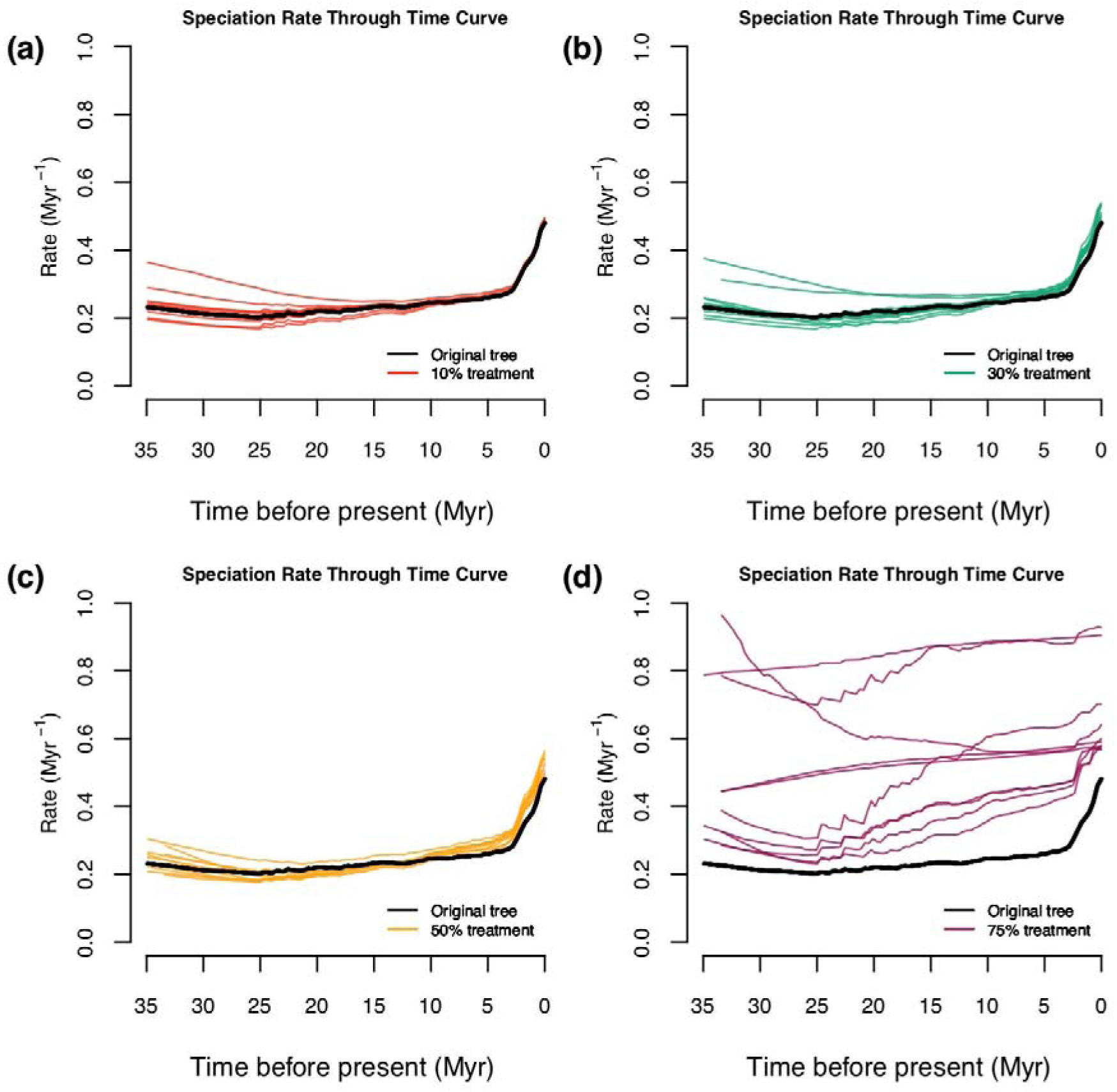
Rate-through-time plots with the random sampling simulations. The thick black line stands for the original Cucurbitaceae 528-tip tree; the color-coded rate-through-time curves were generated by 10 random trees each under 10%, 30%, 50%, and 75% of taxa randomly dropped. The results for all sampling treatments were very similar to the full empirical sampling result except for the most extreme dropping experiment (75% of tips).

*DR—*In contrast to RPANDA and BAMM, DR rates decreased with decreasing sampling effort from 0.3599 Myr^-1^ (10% random drop) to 0.1910 Myr^-1^ (75% random drop; Fig. 5d; Appendix S2.4). The DR rates were significantly different across all treatment comparisons (Tukey HSD; see Appendix S2.5).

*Summary—*As observed with empirical sampling among the 17 rosid orders (above), the estimated contemporary speciation rates increased in RPANDA and BAMM with decreasing sampling effort (10% to 75% random drop; Fig. 5a,c), while rates estimated in DR decreased with decreased sampling (Fig. 5d).

#### Cucurbitaceae test case—Random sampling simulation with backbone taxonomic addition

*RPANDA—*Under random sampling with addition of backbone taxa, the estimated tip speciation rate decreased with decreasing sampling effort (in contrast to random sampling alone; see above), ranging about four-fold from 0.3740 Myr^-1^ (10% backbone-addition; comparable to the 10% random drop treatment, above) to 0.0966 Myr^-1^ (75% backbone-addition; Appendix S2.4). The 10% backbone-addition treatment was significantly higher in contemporary speciation rate than all other treatments (Fig. 5e); no other treatment comparisons were significant (Tukey HSD; see Appendix S2.6).

*BAMM—*As with RPANDA, estimated mean tip speciation rates decreased with decreasing sampling effort and backbone-addition, although the effect was smaller, ranging from 0.4054 (10% random drop & add in) Myr^-1^ to 0.3412 (75% random drop & add in; Fig. 5g; Appendix S2.4). The 10% backbone-addition treatment was significantly higher in contemporary speciation rates than all other treatments; the remaining treatment comparisons were not significant (Tukey HSD; see Appendix S2.6).

Unlike tip speciation rates, decreasing sampling effort with backbone-addition resulted in increased estimated tree-wide speciation rates, ranging from 0.3871 Myr^-1^ (10% random drop & add in) to 0.9545 Myr^-1^ (75% random drop & add in; Fig. 5f; Appendix S2.4). In this case, the tree-wide rates were higher than the tip rates, indicating that the sampling scenario induced early-burst inferences (below). The 10% backbone-addition treatment was significantly lower in contemporary speciation rates than all other treatments; no other treatment comparisons were significant (Tukey HSD; see Appendix S2.6).

Rate-through-time plots from these backbone-addition trees all show a similar pattern of inferring spurious early bursts of diversification (Fig. 7) that were not reconstructed in the original Cucurbitaceae tree (Fig. 7; black curve). Unsurprisingly, these bursts correspond to nodes where backbone taxonomic data were added in these trees.

**Fig. 7.**
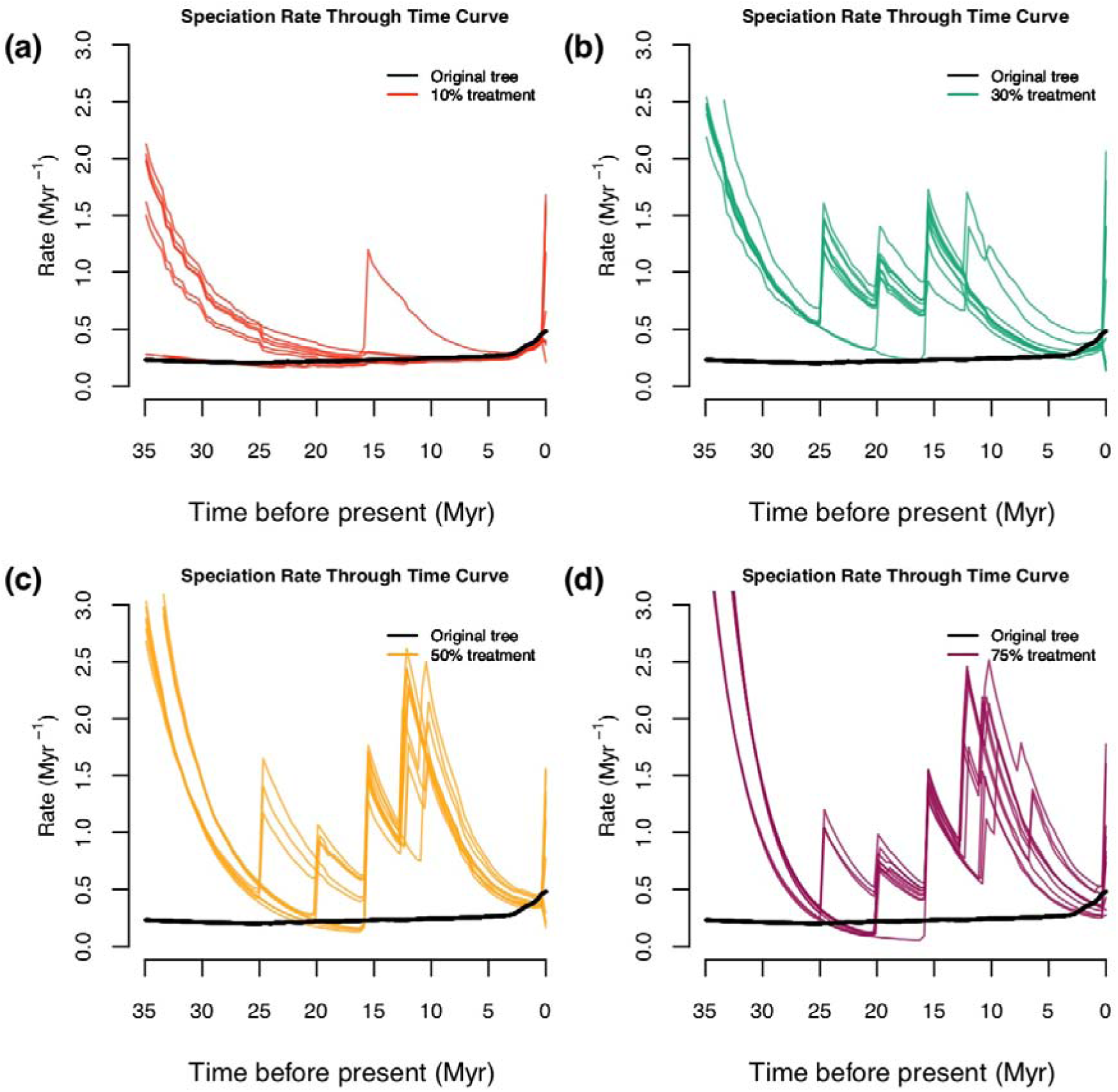
Rate-through-time plots with the random sampling simulations with backbone-addition. The thick black line stands for the original Cucurbitaceae 528-tip tree; the color-coded rate-through-time curves were generated by 10 random trees each under 10%, 30%, 50%, and 75% of taxa randomly dropped and added in as backbone taxonomic data. With moderate missing taxa (10% dropped), few spurious early bursts were inferred, but these were frequent with more missing taxa.

*DR—*DR rates decreased with decreasing sampling effort from 0.3372 Myr^-1^ (10% random drop & add in) to 0.1397 Myr^-1^ (75% random drop & add in; Fig. 5h; Appendix S2.4). The DR rates estimated from all four-level backbone-addition treatments were significantly different for all group comparisons (Tukey HSD; see Appendix S2.6).

*Summary—*Using backbone taxonomic addition to account for missing taxa did not prevent under- or overestimated tip speciation rates. Adding backbone taxa tended to result in the inference of spurious early bursts of diversification (Fig. 7), consistent with the empirical results for the 100k-tip tree (above).

#### Cucurbitaceae test case—Representative sampling simulation

*RPANDA—*Under a representative sampling scenario, the mean tip speciation rate for representative sampling simulations was 0.3022 Myr^-1^ (Fig. 8; see Appendix S2.7), lower by ∼1.5 fold than that for the complete Cucurbitaceae tree (0.4635 Myr^-1^); hence, estimated speciation rates decreased with decreased sampling, opposite the pattern recovered above with random sampling but similar to that recovered with random sampling with backbone-addition.

**Fig. 8.**
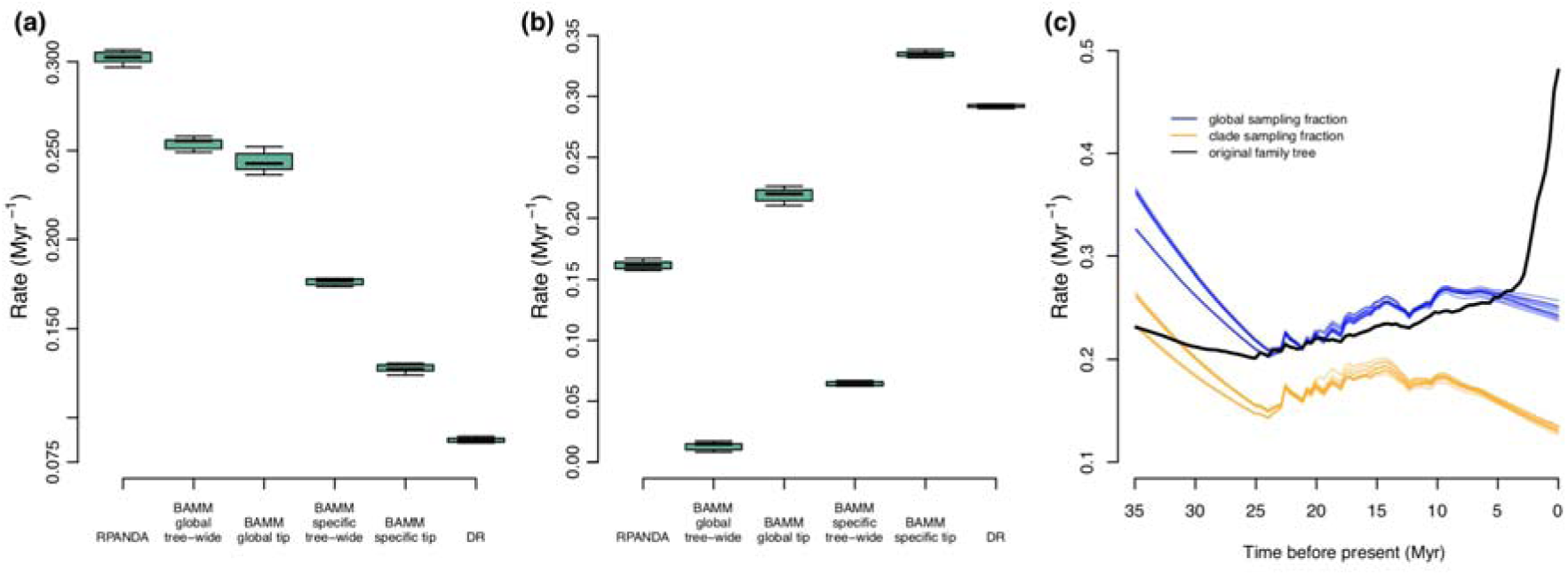
Comparisons of tip speciation rate for full empirical and representative sampling levels for RPANDA, BAMM, and DR using Cucurbitaceae data. (a) Boxplot of contemporary speciation rate and tree-wide rate (BAMM) of the 10 random genus-level tree results estimated by RPANDA, BAMM, and DR. (b) Boxplot showing rate differences by subtracting rates in (a) from those inferred from the family-level 528-tip tree; 0 would indicate identical results. Note that in some cases the magnitude of the difference is nearly as large as the overall speciation rate. (c) Color-coded rate through time plots in BAMM showing rate differences among global sampling fraction (blue), clade-specific sampling fraction (orange), and original family tree (black). Abbreviations for boxplot figures: BAMMglobal tip = tip speciation rates estimated with global sampling fractions; BAMMglobal tree-wide = tree-wide speciation rates estimated with global sampling fractions; BAMMspecific tip = tip speciation rates estimated with clade-specific sampling fractions; BAMMspecific tree-wide = tree-wide speciation rates estimated with clade-specific sampling fractions.

*BAMM—*Unlike RPANDA, BAMM has two approaches for handling incomplete sampling, both implemented here: specifying either clade-specific or global missing taxon parameters. While global sampling fractions were used elsewhere, we included clade-specific sampling fractions here to match common methods used for family-level trees and other backbone phylogenetic data. In the global sampling fraction scenario, mean tip speciation rates (0.1275 Myr^-1^) were lower than the global tree (0.4625 Myr^-1^) while mean tree-wide speciation rates (0.2539 Myr^-1^) were higher than the global tree (0.2408 Myr^-1^). Clade-specific sampling fractions resulted in unilaterally lower estimated speciation rates; both mean tip rates (0.1275 Myr^-1^) and mean tree-wide speciation rates (0.1764 Myr^-1^) were lower than those estimated from the global tree (0.4625 Myr^-1^ and 0.2408 Myr^-1^, respectively; Fig. 8; Appendix S2.7).

Rate-through-time plots (Fig. 8c) were similar to the mean rate results. Global sampling factions tended to increase the scaling of the entire rate curve, with up to ∼two-fold higher speciation rates (at the present), compared to assigning cladewise sampling fractions; the global sampling fraction result was closer to the total Cucurbitaceae tree. While the scaling was different, the rate through time curves were similar in completely failing to detect the burst of speciation rates towards the present seen in the total Cucurbitaceae tree (Fig. 8c); instead, BAMM inferred a spurious early burst of speciation rates at the root (see also backbone-addition, above).

*DR—*The mean DR tip rate for the representative sampling trees was 0.0875 Myr^-1^, far lower than the total Cucurbitaceae tree (0.3794 Myr^-1^), as well as lower than the other rates estimated by RPANDA and BAMM (Fig. 8a-b).

*Summary—*Across methods, representative sampling results in lower tip speciation rate estimates and similar to backbone-addition (above), consistent with these results being driven solely by a failure to sample nodes. However, tree-wide speciation rates were higher on average; rate through time curves (Fig. 8c) showed that this behavior is due to failure to detect recent bursts of speciation and instead inferring higher rates of evolution at earlier time intervals (see also Cusimano et al., 2010).

## IV. Discussion

We found surprisingly diverse effects of sampling effort on inferences of diversification using the methods we employed. Overall, BAMM showed the greatest robustness to incomplete sampling. In BAMM, all random taxon-dropping treatments resulted in statistically identical tip speciation rates with the exception of the most extreme treatment (dropping 75% of taxa; Fig. 5b-c), where estimated tip speciation rate increased dramatically (Appendix S2.4). BAMM also tended to be more robust to the other sampling scenarios, with the exception of representative sampling, where no method was robust. Tree-wide speciation rates and rate-through-time curves in BAMM showed similar patterns (Figs. 6-7), although in some cases these metrics were more sensitive to incomplete sampling than tip speciation rates.

In contrast to BAMM, both RPANDA and DR were highly sensitive to missing taxa. For most analyses, the effect of all incomplete sampling scenarios using RPANDA and DR was disturbingly near-linear (e.g., Fig. 5a, d), in contrast to the threshold behavior of BAMM. Methods also differed in the direction of parameter bias in response to incomplete sampling; DR in all cases resulted in underestimates of tip speciation rates, while BAMM and RPANDA under- or overestimated speciation rates compared to the complete tree, dependent on sampling scenario.

### Opposing bias patterns in representative and random sampling

Under the random sampling scenarios simulated here, speciation estimates *increased* in both RPANDA and BAMM with decreasing sampling efforts (i.e., they were overestimated; Fig. 5). In contrast, representative sampling resulted in decreased estimates of tip speciation rate across methods. Interestingly, in contrast to random sampling, BAMM tip rates were not robust to representative sampling strategies, and these simulations exhibited some of the highest rate estimate differences from the complete Cucurbitaceae tree (Fig. 8b; Appendix S2.7).

Only BAMM and RPANDA showed differential bias patterns, whereas with DR (which does not model taxon absence), decreased sampling always resulted in underestimates of speciation rates. This suggests that modeling taxon absence can result in an “overcorrection” that overestimates rate parameters, even in our taxon-dropping perturbations that were random and therefore matched modeling assumptions. These results make intuitive sense and to some extent are consistent with previous literature (e.g., Cusimano and Renner, 2010). While we attempted to account for incomplete sampling, typically, missing species must be modeled as randomly missing in most implementations of diversification methods. Representative sampling can be seen as a form of sampling bias in that it selectively preserves long phylogenetic branches while dropping short branches. This will have the effect of masking recent, shallow radiation events and pushing apparent diversification patterns backwards in time and depressing estimates of extinction (see Cusimano and Renner, 2010; Höhna et al., 2011). Rate-through-time plots in BAMM exemplify this effect (Figs. 8c, Appendix S3); representative sampling flattened inferred curves and essentially erased any signal of recent diversification, an effect only seen in random sampling with the most extreme scenario (75%; Fig. 6). Instead of a recent burst, representative sampling tends to result in spurious inferences of early bursts not evident with improved sampling (see also Cusimano and Renner, 2010). Understanding this bias is important, as typical molecular phylogenetic sampling schemes seek to represent deep phylogenetic branches disporportionately (Höhna et al., 2011); hence genetic resources like GenBank are likely to be populated primarily by data from studies that used representative sampling schemes.

*Comparison with an angiosperm-wide study—*As an additional exploration of sampling protocols, our BAMM mean speciation rates for the molecular-only trees (9k-tip and 20-k tip; Appendix S2.1) can be directly compared to a recent angiosperm-wide analysis in BAMM exemplifying very coarse representative sampling (Magallón et al., 2018; cf. Supplementary Data “aob-18219-s06”) covering 792 species or ∼0.2% of angiosperm species richness. While Magallón et al. (2018) accounted for incomplete sampling with similar methods to our current study, the difference in results is remarkable. Our estimates of speciation rate with stronger sampling in the same rosid orders (including tree-wide averages and rate-through-time plots) were uniformly higher, the difference sometimes exceeding an order of magnitude (e.g., compare Sapindales, Myrtales, and Vitales; Fig. 3 in Magallón et al., 2018). The mean clade speciation rates we obtained from BAMM ranged up to ∼2.5 Myr^-1^ for the 9k-tip tree and ∼1.7 Myr^-1^ for the 20-tip tree, all values consistent with other rapidly diversifying plant taxa (scaling of plant diversification rates is reviewed in Lagomarsino et al., 2016). All mean clade speciation rates reported in Magallón et al. (2018) were at least 5-fold smaller in magnitude, and even the highest individual lineage speciation rates were at least 2-fold smaller. Unsurprisingly, this angiosperm backbone tree failed to recover signatures of recent diversification; rate curves (Magallón et al., 2018: Fig. 3) were strongly flattened compared to our results, particularly for rate variation within the last ∼15 million years, consistent with our representative sampling experiments (Figs. 8c, Appendix S3). Previous work using coarse phylogenetic sampling with semiparametric methods (Magallón and Sanderson, 2001) had similar scaling of diversification rates to Magallón et al. (2018). The magnitude of this downscaling of speciation rate likewise is similar to that between our molecular-only trees (9k-tip and 20k-tip) and our tree with added taxonomic backbone data (100k-tip; Appendix S3). These observations, along with our sampling manipulation experiments, suggest caution in interpreting the results from diversification studies sampling a very small proportion of species-level diversity with backbone trees and relying heavily on taxonomic data to cover sampling gaps.

### Impact of backbone taxonomic addition

Thus far, we have focused on our 9k- and 20k-tip trees containing only taxa with molecular data. Diversification patterns observed with the 100k-tip tree using backbone taxonomies were remarkably different across methods; the differences mainly comprised (1) spurious inference of early bursts of speciation and (2) depression or exaggeration of tip speciation rates. This difference was consistent across analyses despite a similar phylogenetic backbone across all trees and a similar overall distribution of clade dates between 100k-tip tree and the 9k- and 20k-tip trees (Fig. 2) without obvious overall bias in node age. Despite considerable interest in using synthetic trees for evolutionary studies, we are aware of no similar studies of the behavior of taxon addition by MRCA, as used in OpenTree (Hinchliff et al., 2015; for alternative probabilistic methods, see Thomas et al., 2013; Rabosky, 2015; Rabosky et al., 2018). Among the three diversification methods we used (RPANDA, BAMM, DR), the 100k-tip tree always resulted in far lower estimated tip speciation rates than observed with the 9k- and 20k-tip trees, usually around 10-fold smaller in magnitude (Figs. 3, Appendix S3; Appendix S2.1-S2.3). Although the magnitude is surprising, this pattern makes intuitive sense given that synthetic phylogenies (100k-tip) were built by insertion of missing taxa at the MRCA of the least inclusive clade of which membership is known (e.g., genus, family, etc.). Assuming correct taxonomic assignments, this approach will result in consistently older node ages than would be inferred with molecular data and an empirical tree (9k- and 20k-tip), pushing back the apparent timing of diversification and therefore depressing estimates of tip speciation rate (Figs. 3c, Appendix S3). Simulating this behavior in our backbone-addition experiments confirmed that this practice results in lower estimates of tip speciation rates (Fig. 5e-h; Appendix S2.4), and rate curves showed that this is largely driven by inferring spurious early bursts of evolution (Figs. 7, Appendix S3). As with the random sampling scenario, tip rates in BAMM were most robust to backbone-addition among the methods employed (Fig. 5g), although overall BAMM rates were very sensitive (Fig. 5f).

## V. Conclusions

We found strong impacts of sampling on diversification inference that were surprisingly diverse, and potentially large enough in magnitude to change evolutionary conclusions. For example, our representative and backbone-addition sampling simulations were sufficient to generate spurious early bursts of speciation and erase signals of recent bursts of speciation. Although improvement of molecular taxon sampling to overcome this heterogeneity would be ideal, for large clades this is not always feasible, necessitating methods that adequately account for missing biodiversity knowledge. Our results indicated greater robustness to moderate incomplete sampling in BAMM, especially for estimating tip speciation rate. Some types of rate metrics were more robust than others and more reliable for poorly sampled datasets; tip speciation rates were generally the most robust.

A frequently used alternative to adding molecular data to a given phylogenetic tree is to incorporate taxonomic knowledge and presumed taxonomic placements, often using backbone addition. To date, the benefits of backbone taxonomic addition (e.g., Jetz et al., 2012; Rabosky et al., 2018; Stein et al., 2018) have largely been assumed rather than demonstrated with test cases. We find here that adding taxa without molecular data has unpredictable effects and was not necessarily more accurate than other approaches. Based on the dramatic inferential differences we observed among analyses, we advise (1) strong caution in the inference of diversification using very poorly sampled trees regardless of method; (2) the use of sensitivity analyses similar to those we implemented in Cucurbitaceae to characterize whether empirical results are conditional on methods that account for missing taxa, and (3) especially strong caution in using summary backbone phylogenies for diversification estimation.

## Acknowledgements

This work was supported by the National Science Foundation (DEB-1208809 to D.E.S.; DBI-1523667 to R.A.F.; DEB-1916632 to R.A.F., R.P.G., P.S.S., and D.E.S), Department of Energy (DE-SC0018247 to R.A.F., R.P.G., P.S.S., and D.E.S.), Dimensions of Biodiversity US-China (DEB-1442280 to P.S.S. and D.E.S.), the Strategic Priority Research Program of the Chinese Academy of Sciences (both XDA19050103 and XDB31000000 to Z.D.C.), and National Natural Science Foundation of China (Grant no. 31590822 to Z.D.C.). We thank the HiPerGator cluster at the University of Florida for providing us extensive computational resources.

## Author contributions

The authors declare no conflict of financial interests. R.A.F. and M.S designed the study; M.A.G. advised with data mining; M.S. conducted the analyses; M.S and R.A.F. conducted data interpretation; M.S., R.A.F., and D.E.S. drafted the manuscript; M.A.G., P.S.S., Z.D.C., and R.P.G. revised the manuscript; D.E.S., P.S.S., R.P.G., and Z.D.C. supervised the work. All authors contributed to and approved the final manuscript.

## Data Accessibility Statement

All results of downstream analyses and R scripts will be uploaded to GitHub upon acceptance.

## SUPPORTING INFORMATION

**Appendix S1 Supplemental methods:**

Appendix S1.1: Diversification analyses implemented by RPANDA

Appendix S1.2: Diversification analyses implemented by BAMM

**Appendix S2 Supplemental Tables:**

Appendix S2.1. Summary table for BAMM analyses (also see Appendix S1.2).

Appendix S2.2. Best models and speciation rates estimated for 9k-, 20k, and 100k-tip trees and each of 17 rosid orders from these trees using RPANDA with nine birth-death models (cf. Appendix S1.1).

Appendix S2.3. Summary table for the DR statistic.

Appendix S2.4. Summary table for diversification simulations in the Cucurbitaceae test case.

Appendix S2.5. Tukey HSD test across the RPANDA, BAMM, and DR methods for the Cucurbitaceae test case under the random taxon-dropping scenario.

Appendix S2.6. Tukey HSD test across the RPANDA, BAMM, and DR methods for the Cucurbitaceae test case under the backbone-addition scenario.

Appendix S2.7. Summary table for diversification analyses for the Cucurbitaceae test case under the representative sampling scenario.

**Appendix S3 Supplemental Figure:** Comparison of rate-through-time plots for each of the 17 rosid orders (a-q).

