## Appendix S1 for "Estimating rates and patterns of diversification with incomplete sampling: A case study in the rosids"

**Appendix S1.1: Diversification analyses implemented by RPANDA**

We used RPANDA v1.4 (Morlon et al., 2016) to fit birth-death models for all the three time-calibrated rosid trees (9k-, 20k- and 100k-tip; Morlon, 2014; Morlon et al., 2016). We tested 9 likelihood-based time-dependent diversification models:

1) Pure birth model, no extinction rate (mu, μ = 0), and constant speciation rate (lambda, λ; hereafter **bcst.d0**)

   # t is time

   # y is a vector of initial values feeding to the functions of λ and μ

   *f.lamb = function(t, y){y[1]}*

   *f.mu = function(t, y){0}*

2) Birth-death model with constant speciation and extinction (here as **bcst.dcst**)

   *f.lamb = function(t, y){y[1]}*

   *f.mu = function(t, y){y[1]}*

3) Pure birth model with exponential variation in speciation rate (here as **bvar.d0**)

   *f.lamb = function(t, y){y[1] * exp(y[2] * t)}*

   *f.mu = function(t, y){0}*

4) Pure birth model with linear variation in speciation rate (here as **bvar.l.d0**)

   *f.lamb = function(t, y){y[1] + y[2] * t}*

   *f.mu = function(t, y){0}*

5) Birth-death model with exponential variation in speciation rate and constant extinction (here as **bvar.dcst**)

   *f.lamb = function(t, y){y[1] * exp(y[2] * t)}*

   *f.mu = function(t, y){y[1]}*

6) Birth-death model with linear variation in speciation rate and constant extinction (here as **bvar.l.dcst**)

*f.lamb = function(t, y){y[1] + y[2] * t}*

*f.mu = function(t, y){y[1]}*

7) Birth-death model with a constant speciation rate and exponential variation in extinction (here as **bcst.dvar**)

*f.lamb = function(t, y){y[1]}*

*f.mu = function(t,y){y[1] * exp(y[2] * t)}*

8) Birth-death model with a constant speciation rate and linear variation in extinction (here as **bcst.dvar.l**)

*f.lamb = function(t, y){y[1]}*

*f.mu = function(t,y){y[1] + y[2] * t}*

9) Birth-death model with exponential variation in speciation and extinction (here as **bvar.dvar)**

*f.lamb = function(t, y){y[1] * exp(y[2] * t)}*

*f.mu = function(t,y){y[1] * exp(y[2] * t)}*

These nine birth-death models were calculated using the function "*fit_bd*" (Morlon et al., 2016). We compared AIC values (Akaike Information Criterion; Akaike, 1974) and calculated Akaike weights (Wagenmakers and Farrell, 2004) to quantify relative model support and choose the best model.

**Appendix S1.2: Diversification analyses implemented by BAMM**

We also performed diversification analyses in BAMM (Bayesian Analysis of Macroevolutionary Mixtures; v2.5.0; Rabosky, 2014) to evaluate speciation rates, examining both tip rates (speciation rates at the present) and tree-wide speciation rates (that is, speciation rates across all tree timeframes including the present), as well as rate-through-time plots. BAMM is able to account for non-random and incomplete sampling, allowing the user to assign tips to clades (e.g., genera in our case) and indicate the total proportion of the clade sampled within the phylogeny.

To examine common alternative practices for accounting for unsampled species in BAMM, we explored the impact of using global sampling probabilities (comprising one parameter of missing species for the entire tree) and species-specific sampling probabilities (comprising missing species parameters for arbitrarily defined clades, often named taxa) on diversification rates implemented in BAMM. For this method, it would require excessive computational resources for BAMM to reach MCMC convergence for the entire 20k-tip and 100k-tip trees. To deal with potential convergence issues, we compared the diversification result from independent analyses of 17 subtrees representing recognized orders recovered in 9k-, 20k-, and 100k-tip rosid trees. We calculated subtree diversification analyses in the same manner for all the 17 rosid orders to render the diversification results comparable. In most cases, four independent MCMC chains of 20,000,000 generations were run for each analysis. The initial rate values and rate shift priors were estimated using the R package "BAMMtools" v2.1.6 (Rabosky, 2014) with the "*setBAMMpriors*" function. For larger rosid subclades, the number of generations in MCMC chains, the number of expected shifts, and rate priors were manually adjusted to ensure MCMC convergence. Parameter effective sample sizes (>200 for both the number of shifts and log likelihoods) and convergence among chains were assessed in the R package “coda” v0.19-1 (Plummer et al., 2006). After removing 10% of the trees as burn-in, we explored the BAMM output using BAMMtools to summarize tip and tree-wide speciation rates. We accounted for recent criticism of BAMM (May and Moore, 2016; Moore et al., 2016; Meyer and Wiens, 2017; Meyer et al., 2018), by comparing BAMM rates with those estimated using RPANDA (Morlon et al., 2016) and DR (Jetz et al., 2012) and using the most updated version of BAMM and BAMMtools (Rabosky et al., 2017), considering the author’s response notes (Rabosky, 2018a,b).
