## Appendix S2 for "Estimating rates and patterns of diversification with incomplete sampling: A case study in the rosids"

**Appendix S2.1.** Summary table for BAMM analyses (also see Appendix S1.2).

| Order | 9k-tip tree | | | | 20k-tip tree | | | | 100k-tip tree | | | |
|---|---|---|---|---|---|---|---|---|---|---|---|---|
| | ES_N_shift | ES_logLik | tree-wide rate | tip rate | ES_N_shift | ES_logLik | tree-wide rate | tip rate | ES_N_shift | ES_logLik | tree-wide rate | tip rate |
| Brassicales | 661.10 | 534.45 | 0.7003 | 0.5881 | 1359.32 | 2430.96 | 0.3953 | 0.7258 | 4579.84 | 20.46 | 0.4221 | 0.0893 |
| Celastrales | 2737.53 | 1967.85 | 0.6262 | 0.7583 | 588.06 | 490.94 | 0.2910 | 0.6069 | 13493.91 | 227.97 | 0.3116 | 0.1369 |
| Crossosomatales | 371.53 | 313.11 | 0.1722 | 0.2329 | 781.13 | 494.86 | 0.1771 | 0.2153 | 3550.46 | 2256.42 | 0.0575 | 0.1481 |
| Cucurbitales | 2507.29 | 2146.06 | 0.3319 | 0.5083 | 600.01 | 230.90 | 0.1576 | 0.4176 | 3859.70 | 204.98 | 0.3308 | 0.0680 |
| Fabales | 217.63 | 321.51 | 0.7193 | 1.0854 | 851.79 | 399.77 | 0.4219 | 1.0855 | 22.90 | 5.74 | 0.9133 | 0.0619 |
| Fagales | 37614.85 | 946.02 | 2.4111 | 2.4074 | 469.09 | 266.18 | 1.6606 | 1.4678 | 407.51 | 1892.23 | 0.3297 | 0.1291 |
| Geraniales | 278.96 | 203.93 | 0.4018 | 0.6493 | 395.97 | 241.41 | 0.5363 | 0.6265 | 1484.07 | 429.74 | 0.3165 | 0.1272 |
| Huerteales | 901.00 | 901.00 | 0.0668 | 0.0514 | 782.36 | 995.11 | 0.0759 | 0.0497 | 8418.80 | 9001.00 | 0.0451 | 0.3934 |
| Malpighiales | 214.93 | 347.15 | 0.5830 | 0.7354 | 784.08 | 2226.57 | 0.4140 | 0.7197 | 139.44 | 4.75 | 0.3832 | 0.0789 |
| Malvales | 298.29 | 349.14 | 0.7598 | 1.2517 | 2946.91 | 815.99 | 0.4837 | 1.2493 | 10250.54 | 4915.10 | 0.3104 | 0.1140 |
| Myrtales | 891.38 | 599.38 | 1.0880 | 1.3626 | 2015.80 | 1439.91 | 1.0803 | 1.6819 | 4.84 | 2.77 | 0.2143 | 0.0433 |
| Oxalidales | 257011.88 | 2615.28 | 0.8868 | 0.9800 | 552.70 | 595.89 | 0.8116 | 0.8827 | 523.91 | 1024.82 | 0.3593 | 0.1454 |
| Picramniales | 782.52 | 901.00 | 0.2804 | 0.2911 | 520.72 | 251.23 | 0.2743 | 0.2846 | 90224.85 | 1843.17 | 0.3092 | 0.0569 |
| Rosales | 400.78 | 343.11 | 2.4848 | 3.2572 | 901.00 | 901.00 | 1.6255 | 2.4513 | 376.55 | 8.51 | 0.4346 | 0.0614 |
| Sapindales | 685.37 | 732.94 | 0.4323 | 0.6527 | 560.40 | 815.48 | 0.1500 | 0.4871 | 7874.50 | 106.53 | 0.2942 | 0.0890 |
| Vitales | 901.00 | 217.16 | 1.1109 | 1.1640 | 272.24 | 378.22 | 0.7648 | 0.9796 | 7046.82 | 1646.90 | 1.1542 | 0.0561 |
| Zygophyllales | 901.00 | 652.91 | 0.2535 | 0.2831 | 901.00 | 770.95 | 0.2026 | 0.2164 | 389.49 | 229.85 | 0.4687 | 0.1325 |
| global tree (mean value) | / | / | 0.7829 | 1.1527 | / | / | 0.5601 | 1.0731 | / | / | 0.3914 | 0.1136 |

Notes:

“ES_N_shift” and “ES_logLik” mean the number of shifts (ES_N_shift) and the log-Likelihood (ES_logLik); two values used to assess post-burn-in effective sample sizes (>200) and convergence among chains (BAMM manual).

Tree-wide rate is the speciation rate across all tree timeframes including the present, obtained from the rate-through-time matrix; unit is $Myr^{-1}$.

Tip rate estimated from the "*getTipRates*" function in BAMMtools; unit is $Myr^{-1}$.

BAMM analyses for the 100k-tip tree, for which 6 orders (Brassicales, Fabales, Malpighiales, Myrtales, Rosales, and Sapindales) could not reach suitable effective sample sizes despite runs in some cases exceeding 400 million generations.

**Appendix S2.2.** Best models and speciation rates estimated for 9k-, 20k, and 100k-tip trees and each of 17 rosid orders from these trees using RPANDA with nine birth-death models (cf. Appendix S1.1).

| Order | 9k-tip tree | | | | 20k-tip tree | | | | 100k-tip tree | | | |
|---|---|---|---|---|---|---|---|---|---|---|---|---|
| | **Model** | **λ** | **AICc** | **AW** | **Model** | **λ** | **AICc** | **AW** | **Model** | **λ** | **AICc** | **AW** |
| Brassicales | bvar.dcst | 0.7079172 | 3688.8008 | 0.21165 | bvar.dvar | 0.9421 | 10307.214 | 0.80624 | bvar.l.d0 | 0.0470732 | 43736.32198 | 0.73157 |
| Celastrales | bvar.dcst | 0.8434964 | 1355.5334 | 0.27767 | bvar.dcst | 0.7436 | 1808.7929 | 0.28739 | bvar.l.d0 | 0.0521688 | 11284.71122 | 0.7319 |
| Crossosomatales | bcst.dcst | 0.3074763 | 185.80348 | 0.4336 | bcst.dcst | 0.2968 | 186.51977 | 0.42584 | bvar.l.d0 | 0.1289888 | 532.7036932 | 0.35819 |
| Cucurbitales | bvar.dvar | 0.623294 | 2945.1235 | 0.99423 | bvar.dvar | 0.5014 | 5275.453 | 0.8382 | bvar.l.dcst | 0.0295644 | 23178.99171 | 1 |
| Fabales | bcst.dcst | 1.3051584 | 14884.703 | 0.37936 | bcst.dcst | 1.2903 | 31594.913 | 0.46384 | bvar.d0 | 0.0414453 | 200995.2753 | 0.41474 |
| Fagales | bcst.dcst | 2.5222767 | 1581.8546 | 0.44699 | bcst.dcst | 2.026 | 2539.5318 | 0.47781 | bvar.l.d0 | 0.0474186 | 15247.38435 | 0.73163 |
| Geraniales | bcst.dcst | 1.1756385 | 853.5773 | 0.42802 | bvar.dvar | 0.8114 | 1725.0176 | 0.4725 | bvar.l.d0 | 0.0965253 | 7219.136959 | 1 |
| Huerteales | bcst.d0 | 0.0340313 | 53.341706 | 0.66871 | bcst.d0 | 0.0364 | 52.481626 | 0.58234 | bcst.dvar.l | 0.2649092 | 174.6549247 | 0.50422 |
| Malpighiales | bvar.dvar | 0.9434892 | 13935.882 | 0.28884 | bvar.dcst | 0.9758 | 24684.048 | 0.21372 | bvar.l.d0 | 0.0182802 | 157435.2067 | 0.73093 |
| Malvales | bvar.dcst | 1.4992228 | 5011.1028 | 0.28675 | bvar.dcst | 1.3722 | 7881.9606 | 0.32356 | bvar.l.d0 | -0.001315 | 49127.59102 | 0.73125 |
| Myrtales | bcst.dcst | 2.3182921 | 4809.0756 | 0.24044 | bvar.dcst | 2.4124 | 8268.0079 | 0.29772 | bvar.l.dcst | -0.006568 | 116233.782 | 1 |
| Oxalidales | bcst.dvar.l | 1.4855975 | 753.60331 | 0.53209 | bcst.dvar.l | 1.0214 | 1388.6102 | 0.37694 | bvar.l.d0 | 0.0497701 | 16649.34027 | 0.73159 |
| Picramniales | bcst.d0 | 0.0771328 | 33.297635 | 0.72901 | bcst.d0 | 0.0765 | 33.33898 | 0.73159 | bvar.l.dcst | 0.0372405 | 370.5424775 | 0.99659 |
| Rosales | bvar.dvar | 4.6098893 | 4950.3889 | 0.95931 | bcst.dcst | 3.056 | 9485.4273 | 0.44212 | bvar.dvar | 0.0258025 | 116962.3346 | 1 |
| Sapindales | bvar.dvar | 0.6382086 | 5733.1687 | 0.5147 | bvar.dvar | 0.5313 | 8847.7864 | 0.92422 | bvar.l.d0 | 0.0269303 | 41719.72204 | 0.73135 |
| Zygophyllales | bcst.dcst | 0.4179997 | 385.42321 | 0.4215 | bcst.dcst | 0.2635 | 582.47463 | 0.40309 | bvar.l.d0 | 0.0257752 | 2800.561744 | 0.72948 |
| Vitales | bvar.dvar | 11.214279 | 335.58215 | 0.99296 | bcst.dcst | 1.5141 | 832.30451 | 0.27539 | bvar.dvar | 0.0188394 | 9630.261453 | 0.83137 |
| global tree | bvar.dvar | 1.3904588 | 62217.036 | 0.38514 | bvar.dvar | 1.3058 | 116622.39 | 0.50235 | bvar.l.d0 | 0.0445547 | 857591.7899 | 0.73367 |

Notes: Model: see Appendix S1.1 for further information.

AICc: the corrected Akaike Information Criterion (AICc) for the fitted model.

AW is Akaike weight (Wagenmakers and Farrell, 2004).

Lambda is the speciation rate (λ) parameter estimated by the fitted birth-death model at the present, see *f.lamb* function in Morlon et al.

(2014); unit is $Myr^{-1}$.

**Appendix S2.3. Summary table for the DR statistic.**

Mean DR tip rates were averaged across the 17 orders and the whole of 9k-, 20k-, and 100k-tip rosid trees, respectively; unit is $Myr^{-1}$.

| **Order** | **9k-tip tree** | **20k-tip tree** | **100-k tip tree** |
|---|---|---|---|
| Brassicales | 0.1216 | 0.5058 | 0.1125 |
| Celastrales | 0.1854 | 0.2210 | 0.0903 |
| Crossosomatales | 0.0840 | 0.0764 | 0.1005 |
| Cucurbitales | 0.2159 | 0.3055 | 0.0888 |
| Fabales | 0.1778 | 0.5975 | 0.1080 |
| Fagales | 0.3241 | 0.5777 | 0.0808 |
| Geraniales | 0.1538 | 0.3214 | 0.0915 |
| Huerteales | 0.0216 | 0.0222 | 0.1133 |
| Malpighiales | 0.1284 | 0.3435 | 0.0931 |
| Malvales | 0.2466 | 0.4294 | 0.1129 |
| Myrtales | 0.1656 | 0.4217 | 0.0727 |
| Oxalidales | 0.0894 | 0.1854 | 0.1011 |
| Picramniales | 0.0303 | 0.0312 | 0.0238 |
| Rosales | 0.3459 | 0.6392 | 0.0599 |
| Sapindales | 0.1910 | 0.2554 | 0.0804 |
| Vitales | 0.1126 | 0.5672 | 0.0450 |
| Zygophyllales | 0.0555 | 0.0697 | 0.0673 |
| global tree (mean value) | 0.1889 | 0.4644 | 0.0902 |

**Appendix S2.4. Summary table for diversification simulations in the Cucurbitaceae test case.**

| | Treatment | Tip no. | RPANDA | | | | Mean DR | BAMM | |
|---|---|---|---|---|---|---|---|---|---|
| | | | model | λ | AICc | AW | | mean tree-wide rate | mean tip rate |
| | Cucurbitaceae_20k_family | 528 | bcst.dcst | 0.4635 | 2868.24 | 0.24 | 0.3794 | 0.2408 | 0.4625 |
| | Cucurbitaceae_20k_genus | 123 | bvar.d0 | 0.2687 | 751.081 | 0.33 | 0.1052 | 0.2571 | 0.2543 |
| random drop | Cucurbitaceae_20k_10% | 475 | bvar.dcst | 0.4688 | 2595.68 | 1.00 | 0.3599 | 0.2466 | 0.4658 |
| random drop | Cucurbitaceae_20k_30% | 370 | bcst.dcst | 0.5471 | 2076.03 | 1.00 | 0.3282 | 0.2516 | 0.4951 |
| random drop | Cucurbitaceae_20k_50% | 264 | bcst.dcst | 0.6283 | 1539.34 | 0.93 | 0.2731 | 0.2491 | 0.5010 |
| random drop | Cucurbitaceae_20k_75% | 132 | bcst.dcst | 0.7264 | 822.306 | 0.51 | 0.1910 | 0.5261 | 0.6508 |
| backbone addition | Cucurbitaceae_20k_10% | 528 | bcst.dcst | 0.3800 | 3016 | 1.00 | 0.3373 | 0.3871 | 0.4054 |
| backbone addition | Cucurbitaceae_20k_30% | 528 | bvar.dvar | 0.1729 | 3297 | 0.96 | 0.2647 | 0.8536 | 0.3662 |
| backbone addition | Cucurbitaceae_20k_50% | 528 | bvar.dvar | 0.1571 | 3382 | 1.00 | 0.2013 | 0.9608 | 0.3661 |
| backbone addition | Cucurbitaceae_20k_75% | 528 | bvar.dvar | 0.0966 | 3276 | 1.00 | 0.1397 | 0.9545 | 0.3412 |

**Appendix S2.5. Tukey HSD test across the RPANDA, BAMM, and DR methods for the Cucurbitaceae test case under the random taxon-dropping scenario.** p.adj means adjusted *p*-value. Values in boldface are significant.

RPANDA

| treatment | diff | lwr | upr | p.adj |
|---|---|---|---|---|
| Cucurbitaceae 20k 30%—10% | 0.0754 | -0.0407 | 0.1916 | 0.31398 |
| Cucurbitaceae 20k 50%—10% | 0.1288 | 0.0127 | 0.2450 | 0.02482 |
| Cucurbitaceae 20k 75%—10% | 0.3644 | 0.2483 | 0.4806 | **2.70e-09** |
| Cucurbitaceae 20k 50%—30% | 0.0534 | -0.0627 | 0.1695 | 0.60715 |
| Cucurbitaceae 20k 75%—30% | 0.2890 | 0.1728 | 0.4051 | **4.77e-07** |
| Cucurbitaceae 20k 75%—50% | 0.2356 | 0.1194 | 0.3517 | **2.10e-05** |

BAMM tip rate

| treatment | diff | lwr | upr | p.adj |
|---|---|---|---|---|
| Cucurbitaceae 20k 30%—10% | 0.0292 | 0.0029 | 0.0556 | 0.022429 |
| Cucurbitaceae 20k 50%—10% | 0.0352 | 0.0060 | 0.0643 | 0.010474 |
| Cucurbitaceae 20k 75%—10% | 0.1850 | 0.1476 | 0.2223 | **0** |
| Cucurbitaceae 20k 50%—30% | 0.0059 | -0.0247 | 0.0365 | 0.959804 |
| Cucurbitaceae 20k 75%—30% | 0.1557 | 0.1172 | 0.1942 | **0** |
| Cucurbitaceae 20k 75%—50% | 0.1498 | 0.1093 | 0.1903 | **0** |

BAMM tree-wide rate

| treatment | diff | lwr | upr | p.adj |
|---|---|---|---|---|
| Cucurbitaceae 20k 30%—10% | 0.0050 | -0.1111 | 0.1210 | 0.999446314 |
| Cucurbitaceae 20k 50%—10% | 0.0024 | -0.1136 | 0.1185 | 0.999933504 |
| Cucurbitaceae 20k 75%—10% | 0.2794 | 0.1634 | 0.3955 | **9.22e-07** |
| Cucurbitaceae 20k 50%—30% | -0.0025 | -0.1186 | 0.1136 | 0.999927428 |
| Cucurbitaceae 20k 75%—30% | 0.2745 | 0.1584 | 0.3906 | **1.31e-06** |
| Cucurbitaceae 20k 75%—50% | 0.2770 | 0.1609 | 0.3931 | **1.10e-06** |

DR

| treatment | diff | lwr | upr | p.adj |
|---|---|---|---|---|
| Cucurbitaceae 20k 30%—10% | -0.0317 | -0.0497 | -0.0137 | **0.00019** |
| Cucurbitaceae 20k 50%—10% | -0.0868 | -0.1048 | -0.0688 | **0** |
| Cucurbitaceae 20k 75%—10% | -0.1689 | -0.1869 | -0.1509 | **0.00e+00** |
| Cucurbitaceae 20k 50%—30% | -0.0551 | -0.0731 | -0.0371 | **4.75e-09** |

| | | | | |
|---|---|---|---|---|
| Cucurbitaceae 20k 75%—30% | -0.1372 | -0.1552 | -0.1192 | **0.00e+00** |
| Cucurbitaceae 20k 75%—50% | -0.0821 | -0.1001 | -0.0641 | **0.00e+00** |

**Appendix S2.6. Tukey HSD test across the RPANDA, BAMM, and DR methods for the Cucurbitaceae test case under the backbone-addition scenario.**

p.adj means adjusted *p*-value. Values in boldface are significant.

RPANDA

| treatment | diff | lwr | upr | p.adj |
|---|---|---|---|---|
| Cucurbitaceae 20k 30%—10% | -0.1220 | -0.2089 | -0.0352 | **0.00303004** |
| Cucurbitaceae 20k 50%—10% | -0.1993 | -0.2862 | -0.1124 | **2.34e-06** |
| Cucurbitaceae 20k 75%—10% | -0.1951 | -0.2819 | -0.1082 | **3.49e-06** |
| Cucurbitaceae 20k 50%—30% | -0.0773 | -0.1641 | 0.0096 | 0.09615573 |
| Cucurbitaceae 20k 75%—30% | -0.0730 | -0.1599 | 0.0138 | 0.12562993 |
| Cucurbitaceae 20k 75%—50% | 0.0042 | -0.0826 | 0.0911 | 0.99918405 |

BAMM tip rate

| treatment | diff | lwr | upr | p.adj |
|---|---|---|---|---|
| Cucurbitaceae 20k 30%—10% | -0.0392 | -0.0595 | -0.0190 | **3.56e-06** |
| Cucurbitaceae 20k 50%—10% | -0.0393 | -0.0595 | -0.0191 | **3.47e-06** |
| Cucurbitaceae 20k 75%—10% | -0.0642 | -0.0844 | -0.0440 | **0** |
| Cucurbitaceae 20k 50%—30% | 0.0000 | -0.0203 | 0.0202 | 0.999999951 |
| Cucurbitaceae 20k 75%—30% | -0.0249 | -0.0452 | -0.0047 | 0.008276584 |
| Cucurbitaceae 20k 75%—50% | -0.0249 | -0.0451 | -0.0047 | 0.00841872 |

BAMM tree-wide rate

| treatment | diff | lwr | upr | p.adj |
|---|---|---|---|---|
| Cucurbitaceae 20k 30%—10% | 0.4664 | 0.3552 | 0.5777 | **1.06E-12** |
| Cucurbitaceae 20k 50%—10% | 0.5737 | 0.4625 | 0.6849 | **0** |
| Cucurbitaceae 20k 75%—10% | 0.5674 | 0.4562 | 0.6786 | **0** |
| Cucurbitaceae 20k 50%—30% | 0.1073 | -0.0040 | 0.2185 | 0.062090622 |
| Cucurbitaceae 20k 75%—30% | 0.1010 | -0.0103 | 0.2122 | 0.086624821 |
| Cucurbitaceae 20k 75%—50% | -0.0063 | -0.1175 | 0.1049 | 0.998705281 |

DR

| treatment | diff | lwr | upr | p.adj |
|---|---|---|---|---|
| Cucurbitaceae 20k 30%—10% | -0.0725 | -0.0904 | -0.0546 | **3.11e-12** |
| Cucurbitaceae 20k 50%—10% | -0.1359 | -0.1538 | -0.1180 | **0** |
| Cucurbitaceae 20k 75%—10% | -0.1976 | -0.2155 | -0.1797 | **0** |
| Cucurbitaceae 20k 50%—30% | -0.0634 | -0.0813 | -0.0455 | **1.26e-10** |

| | | | | |
|---|---|---|---|---|
| Cucurbitaceae 20k 75%—30% | -0.1251 | -0.1429 | -0.1072 | **0** |
| Cucurbitaceae 20k 75%—50% | -0.0617 | -0.0795 | -0.0438 | **2.57e-10** |

**Appendix S2.7. Summary table for diversification analyses in Cucurbitaceae test case under the representative sampling scenario.**

| genus tree | RPANDA | | | | BAMM | | | | DR |
|---|---|---|---|---|---|---|---|---|---|
| | | | | | global sampling | | species-specific sampling | | |
| | Model | lamda | AICc | AW | mean tree-wide | mean tip rate | mean tree-wide | mean tip rate | mean DR rate |
| Cucurbitaceae 20k genus 1 | bvar.d0 | 0.2970 | 394.14 | 0.3164 | 0.2559 | 0.2469 | 0.1780 | 0.1275 | 0.0883 |
| Cucurbitaceae 20k genus 2 | bvar.d0 | 0.3072 | 390.65 | 0.3188 | 0.2554 | 0.2391 | 0.1774 | 0.1251 | 0.0868 |
| Cucurbitaceae 20k genus 3 | bvar.d0 | 0.3061 | 393.06 | 0.3193 | 0.2509 | 0.2401 | 0.1730 | 0.1266 | 0.0858 |
| Cucurbitaceae 20k genus 4 | bvar.d0 | 0.2998 | 390.48 | 0.2950 | 0.2565 | 0.2494 | 0.1785 | 0.1305 | 0.0877 |
| Cucurbitaceae 20k genus 5 | bvar.d0 | 0.3044 | 395.77 | 0.3145 | 0.2557 | 0.2408 | 0.1781 | 0.1274 | 0.0898 |
| Cucurbitaceae 20k genus 6 | bvar.d0 | 0.3055 | 394.62 | 0.3367 | 0.2511 | 0.2487 | 0.1746 | 0.1307 | 0.0885 |
| Cucurbitaceae 20k genus 7 | bvar.d0 | 0.2998 | 394.99 | 0.3244 | 0.2559 | 0.2445 | 0.1783 | 0.1305 | 0.0891 |
| Cucurbitaceae 20k genus 8 | bvar.d0 | 0.2968 | 394.31 | 0.3151 | 0.2490 | 0.2373 | 0.1751 | 0.1262 | 0.0861 |
| Cucurbitaceae 20k genus 9 | bvar.d0 | 0.3020 | 394.34 | 0.3263 | 0.2593 | 0.2545 | 0.1766 | 0.1272 | 0.0875 |
| Cucurbitaceae 20k genus 10 | bvar.d0 | 0.3030 | 388.85 | 0.2934 | 0.2492 | 0.2354 | 0.1746 | 0.1231 | 0.0858 |
| mean rates | / | 0.3022 | / | / | 0.2539 | 0.2437 | 0.1764 | 0.1275 | 0.0875 |
| Cucurbitaceae family tree | bcst.dcst | 0.4635 | 2868.24 | 0.2406 | 0.2408 | 0.4625 | / | / | 0.3794 |
