## Appendix S3 for "Estimating rates and patterns of diversification with incomplete sampling: A case study in the rosids"

**Appendix S3.** Comparison of rate-through-time plots for each of the 17 rosid orders (a-q). Each panel coplots results from the 9k- (dashed line), 20k- (solid line), and 100k-tip trees (dotted line). For the orders Cucurbitales, Fabales, and Vitales, speciation rates had much higher scaling for the 100k-tip tree; in these cases, thumbnail rate-through-time plots are provided for the 9k- and 20k-tip trees to show further details. These curves show that (1) tree-wide speciation rates were estimated differently based on different sampling scales; (2) sparse sampling tends to generate flattened curves and erase recent diversification signals (9k-tip tree vs. 20k-tip tree); (3) the 100k-tip tree tended to produce lower speciation rates at present (0 Myr); and (4) the 100k-tip tree, where missing taxa are taxonomically assigned to the tree backbone, pushes back the apparent timing of diversification, causing detection of spurious early bursts of evolution.

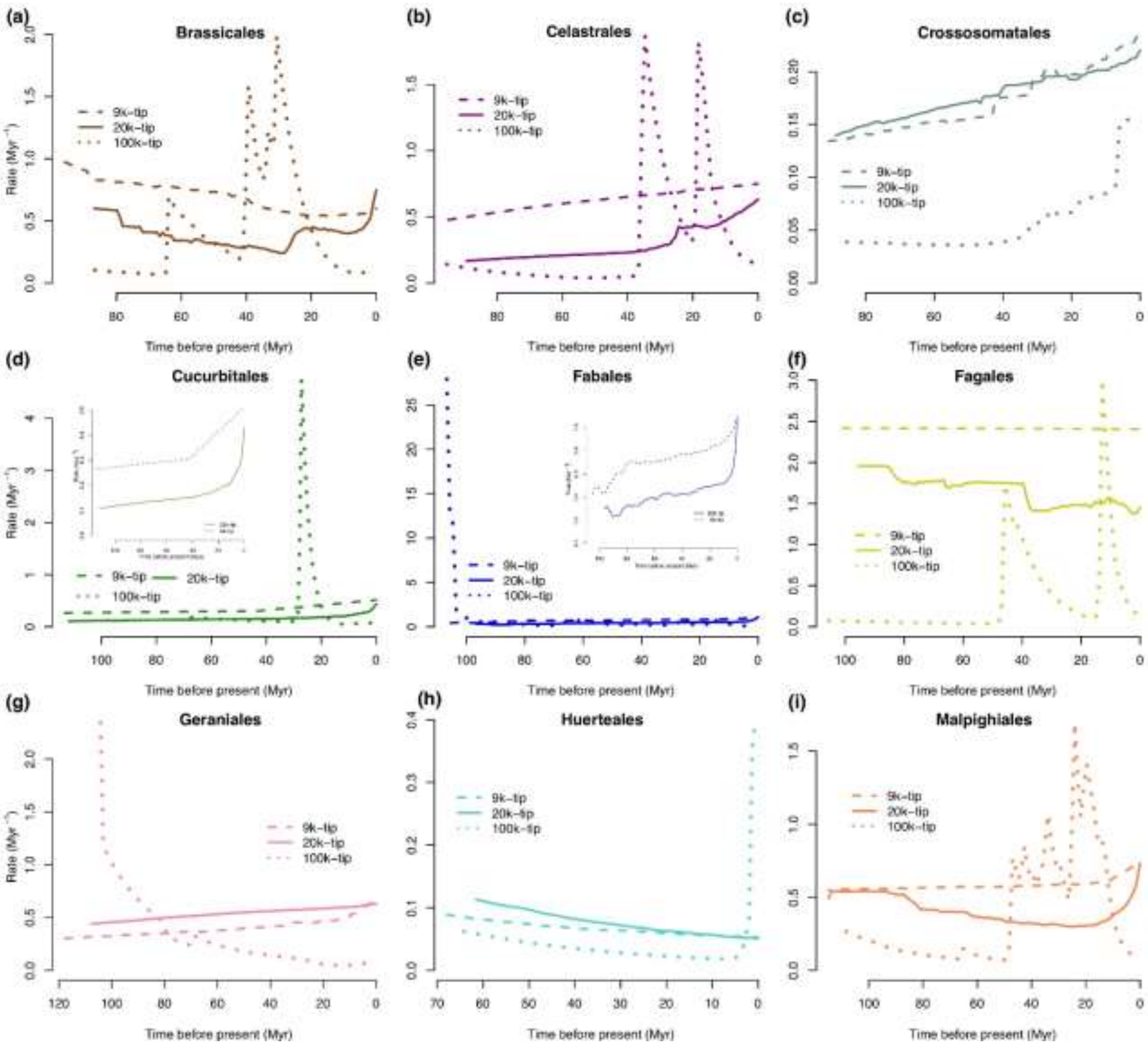
(a)
Brassicales
(b)
Celastrales
(c)
Crossosomatales
(d)
Cucurbitales
(e)
Fabales
(f)
Fagales
(g)
Geraniales
(h)
Huerteales
(i)
Malpighiales
9k–tip
20k–tip
100k–tip
Rate (Myr⁻¹)
Time before present (Myr)

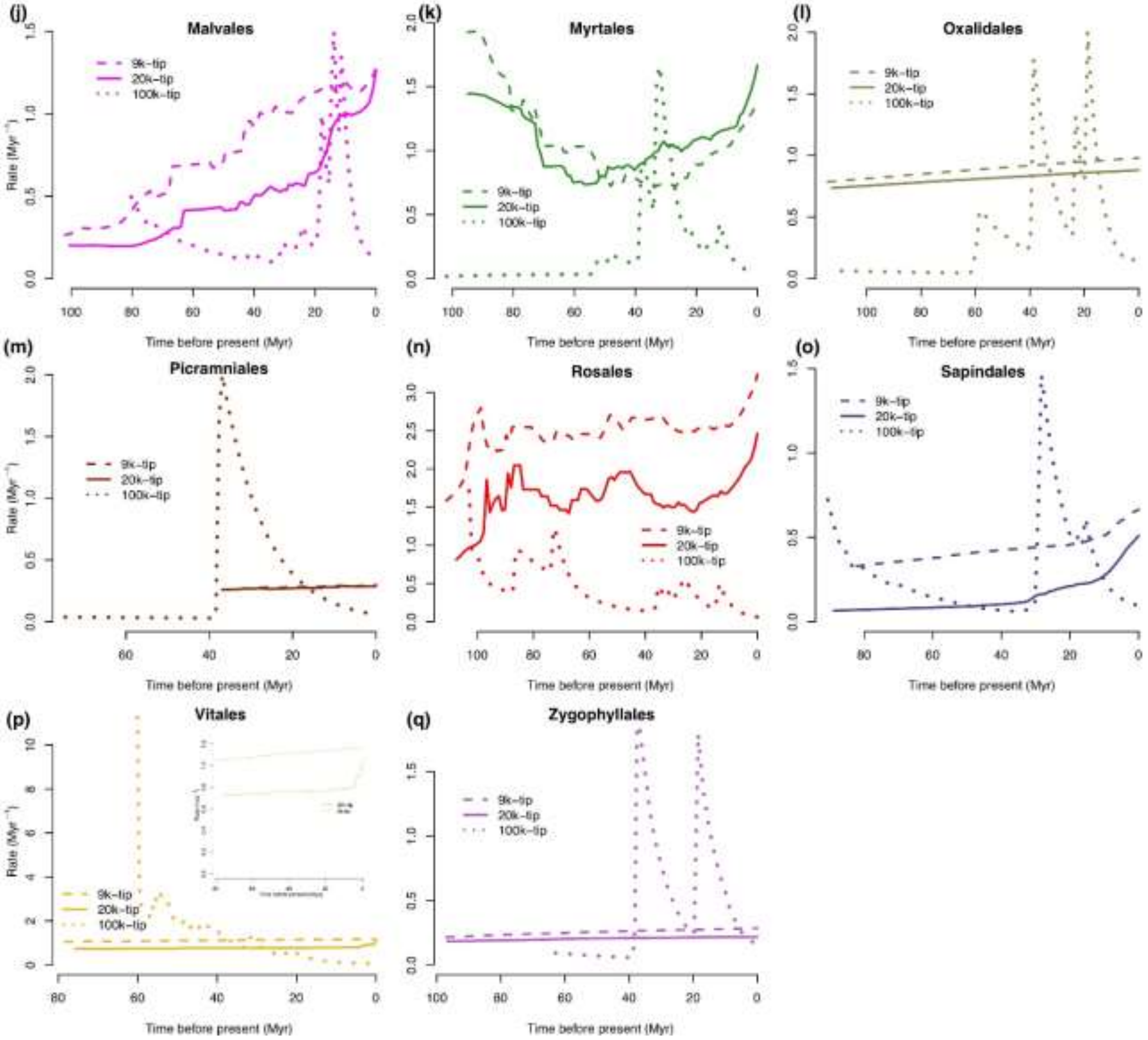
(j)
Malvales
(k)
Myrtales
(l)
Oxalidales
(m)
Picramniales
(n)
Rosales
(o)
Sapindales
(p)
Vitales
(q)
Zygophyllales
9k-tip
20k-tip
100k-tip
Rate (Myr⁻¹)
Time before present (Myr)
